# The RNA Binding proteome of axonal mRNAs in sympathetic neurons

**DOI:** 10.1101/2022.11.23.517728

**Authors:** Raphaëlle Luisier, Catia Andreassi, Antonella Riccio

## Abstract

**Background:** Neurons are morphologically complex cells that rely on the compartmentalization of protein expression to develop and maintain their cytoarchitecture. Targeting of RNA transcripts to axons is one of the mechanisms that allows rapid local translation of proteins in response to extracellular signals. 3’ untranslated regions (UTRs) of mRNA are non-coding sequences that play a critical role in determining transcript localisation and translation by interacting with specific RNA binding proteins (RBPs). However, how 3’UTRs contribute to mRNA metabolism and the nature of RBP complexes responsible for these functions remain elusive.

**Results:** We performed 3’ end sequencing of RNA isolated from axons and cell bodies of sympathetic neurons exposed to either Nerve Growth factor (NGF) or Neurotrophin 3 (NT3). NGF and NT3 are growth factors essential for sympathetic neuron development that act through distinct signalling mechanisms. Whereas NT3 is thought to act only locally, NGF signals back from axons to the cell bodies. We discovered that both NGF and NT3 affect transcription and alternative polyadenylation and induce the localisation of specific 3’UTR isoforms to axons. The finding that many transcripts with short 3’UTR were detected only in axons suggest that these may undergo local post-transcriptional remodelling. The integration of our data with CLIP-sequencing data revealed that long 3’UTR isoforms associate with RBP complexes in the nucleus, and once in axons, regulate cytoplasmic 3’ UTR isoform cleavage into shorter isoform.

**Conclusions:** Our findings shed new light on the complex interplay between nuclear polyadenylation, mRNA localisation and local 3’UTR remodelling in developing neurons.

**Graphical abstract:** 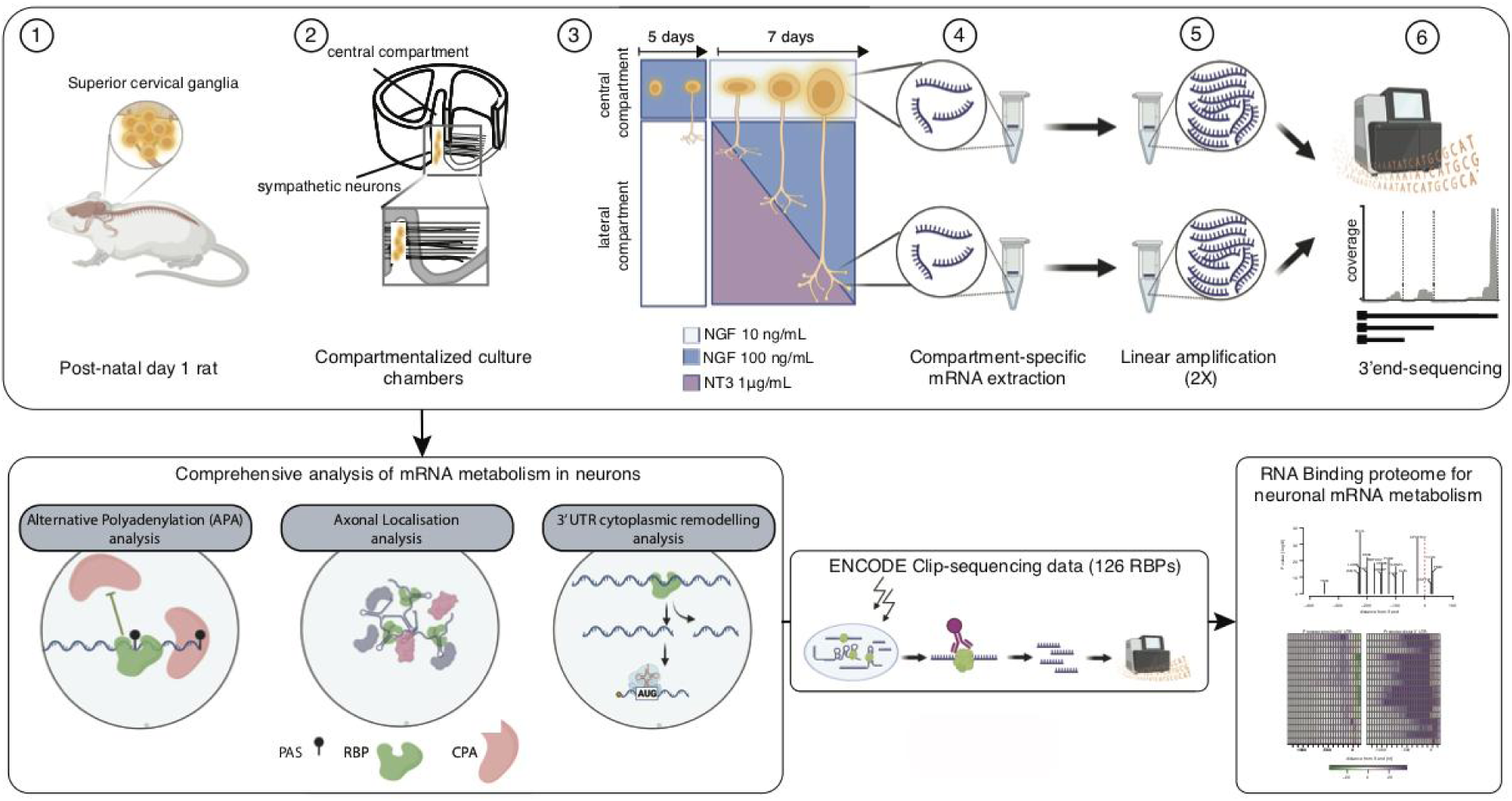

## BACKGROUND

RNA transport to axons and local translation in response to extracellular cues play an essential role in mediating axon growth and cell survival of developing neurons. Although initially the presence of RNA transcripts in axons was considered an oddity, a wealth of studies demonstrated that RNA transport and translation in axons is a widespread phenomenon observed in most neuronal cell types and across the species [1]. We now know that local synthesis of axonal transcripts is essential for many biological functions, including axon extension, growth cone turning and nerve regeneration after injury [2]. During neuronal development, axons extend over long distances to reach their targets, driven by their growth cone, which responds rapidly to local cues along its migratory path [3]. At early developmental stages, sympathetic neurons respond to neurotrophin 3 (NT3) released from the vasculature surrounding the axons [4]. NT3 elicits a local signalling sufficient to support axon growth and cell survival of the relatively immature neurons [5, 6]. However, at later stages, when axons have reached a considerable length, this local mechanism is no longer suitable to propagate the signalling back to the cell bodies in a reliable manner [7]. As distal axons and growth cones approach their final targets, they release Nerve Growth Factor (NGF) that is then internalised within signalling endosomes and transported back to the cell bodies where it activates transcription [6, 8, 9]. Although in this cell system both neurotrophins chiefly bind to the same tyrosine kinase receptor TrkA, their ability to signal to the nucleus has been linked to their differential internalization into signalling endosomes and retrograde transport to cell bodies. Because the NT3-TrkA complexes are not found in signalling endosomes and are not retrogradely transported, NT3 is considered to be a poor activator of gene expression and to act locally in axons [6].

In addition to conveying the genetic data from the chromatin to the translational machinery, mRNA transcripts carry further information stored in their untranslated regions (UTRs). The 3’UTRs play a key role in regulating many aspects of RNA metabolism, including transcript localization, mRNA stability and translation by interacting with RNA binding proteins (RBPs) [10–12]. RBP complexes are initially assembled co-transcriptionally and are essential for regulating mRNA splicing, polyadenylation site (PAS) choice and alternative polyadenylation (APA), and mRNA nuclear export [13, 14]. Most RBPs are multifunctional and RBP complexes undergo extensive remodelling in the cytoplasm. For example, RBPs interaction with elements in the 3’UTR mediate mRNA targeting to dendrites and axons, and this event is necessary for the establishment of synapses, axon growth and neuronal development [15–18]. Critically, incorrect processing and delivery of mRNA causes severe developmental defects and has been implicated in many neurological disorders [19].

We performed 3’end sequencing of RNA localised in axons and cell bodies of sympathetic neurons whose axons were exposed to either NGF or NT3, and found that both neurotrophins induce a robust transcriptional response with many transcripts specifically regulated when either growth factor was applied to the axon terminals. NGF and NT3 also induced Alternative Polyadenylation (APA), thereby generating multiple 3’UTR isoforms. A novel Bayesian procedure developed to accurately identify localised 3’ UTR isoforms revealed hundreds of isoforms that are actively and specifically transported in axons. Notably dozens of isoforms expressing the short 3’UTR were exclusively located in axons, suggesting that 3’UTRs may undergo local post-transcriptional remodelling [20] in response to either NGF or NT3. Integration of our findings with predicted binding events inferred from publicly available CLIP-sequencing data for 126 RNA binding proteins (RBP) revealed that RBP complexes associated with the downstream regions of the short 3’ UTRs compete with cleavage factors and promote the expression of long 3’ UTRs that are transported in axons. Once in the axonal compartment, the complexes induce the cleavage of the long 3’UTR through the interaction with additional RBPs.

## RESULTS

### NGF and NT3 applied to distal axons induce distinct transcriptional programmes

To identify mRNAs localised in sympathetic cell bodies and axons grown in the presence of NGF or NT3, we combined compartmentalized cultures and 3’UTR sequencing [20]. Neonatal rat sympathetic neurons were cultured in chambers that allow the physical separation of cell bodies from distal axons (**Supplementary Fig. 1A**). Sympathetic neurons are especially suitable for compartmentalized cultures because they grow in culture as a highly homogeneous population without glial cells. Neurons were seeded in the central compartment with NGF (100 ng/ml) and after 5 days, the NGF was lowered to 10ng/ml in the cell bodies, and NGF or NT3 were supplied at high concentration to the peripheral compartments to stimulate axon growth. After 7 additional days *in vitro*, RNA was isolated from either cell bodies or axons, subject to 2 rounds of linear amplification and sequenced by using 3’end-Seq [20]. This technique allowed the sequencing of transcripts 3’ends independently of the length of the transcript [20] (see methods for a detailed description of the protocol used; Fig. 1A). Analyses of the reads obtained from two independent biological replicates indicated the high reliability of the RNA sequencing in both compartments (**Supplementary Fig. 1B**). The lower correlation between the replicates from the axonal compartment is likely due to the lower amount of RNA isolated in axons.

**Figure 1.**
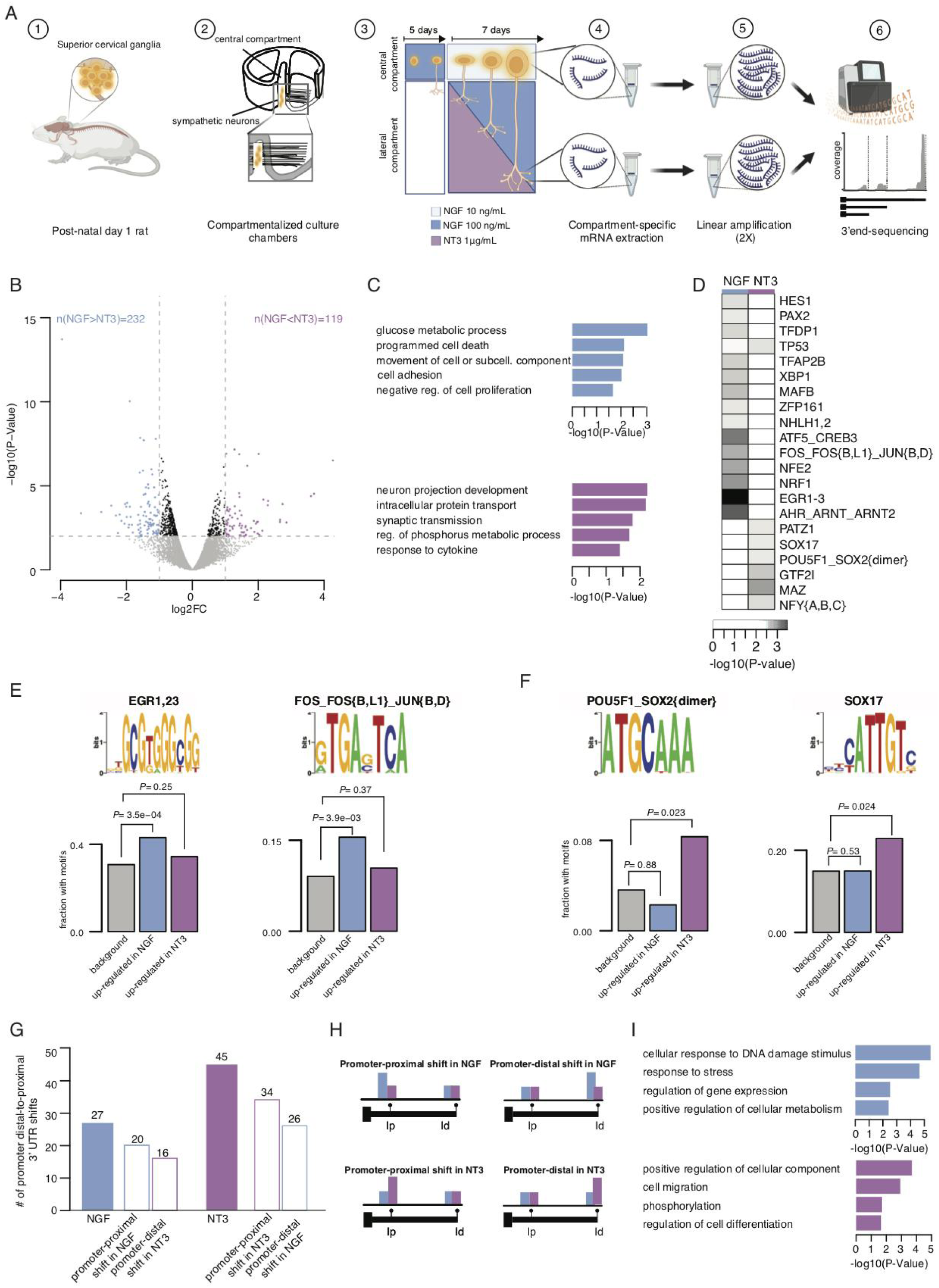
NGF and NT3 are equally capable of regulating transcriptional changes when applied to distal axons. (**A**) Schematic of the experimental set-up. **(B)** Volcano plot representing log2 fold-change (log2FC) in gene expression values between neurons whose axons were exposed to NGF or NT3, and corresponding P-values (-log10). Blue dots = genes significantly up-regulated in NGF; purple dots = genes significantly up-regulated in NT3 (FC>1.5 and P-value <0.01). **(C)** Enrichment scores of GO biological pathways associated with up-regulated genes in NGF- (*upper*) and NT3-treated (*lower*) neurons. **(D)** Fisher enrichments in predicted binding sites for 21 transcription factors in the 1000 nucleotide promoter region of the up-regulated genes in NGF or NT3 treated neurons. **(E,F)** Sequence logos of predicted motifs bound by transcription factors (*upper*) and fractions of promoter regions with these motifs (*lower*) in the total pool of expressed genes (grey bar), the genes up-regulated in NGF (blue bar) and the genes up-regulated in NT3 (purple bar). (Fisher enrichment test). **(G)** Comparison of the total number of proximal-to-distal promoter 3’ UTR shifts in either condition (full bars) and by events subtype, as depicted in (H) underlying these events (empty bars). **(H)** Schematic of relative usage of the promoter-proximal (Ip) and promoter-distal (Id) 3’ UTR isoforms underlying differential 3’ UTR isoform usage between NGF and NT3. **(I)** Enrichment scores of GO biological pathways associated with significant increase in distal-to-proximal promoter 3’ UTR usage in NGF compared to NT3 (*upper*) and vice-versa (*lower*).

We first tested whether exposure of distal axons to either NGF or NT3 leads to transcriptional changes in cell bodies. Differential gene expression analysis identified 232 genes that were up-regulated in NGF treated neurons compared to NT3 (FC>1.5 and P value<0.01, **Fig. 1B**, **Supplementary Table 1**). The genes were enriched in terms related to cell adhesion and glucose metabolism (**Fig. 1C**, *upper*). Conversely, the expression of 119 genes was increased in the cell bodies of neurons whose axons were exposed to NT3, and they were enriched for terms such as neuron projection and synaptic transmission (**Figs. 1C**, *lower*, **Supplementary Table 2**). Transcription factor binding site (TFBS) enrichment analysis showed a neurotrophin-specific transcriptional regulation for these two categories of differentially expressed genes (**Fig. 1D**, **Supplementary Table 3**). The 1,000 nucleotide promoter regions of the genes up-regulated in NGF were enriched in 14 TFBS motifs that included motifs bound by EGR1, EGR2 and EGR3 (P-Value=3.46E-04, 43% of the promoter regions of NGF up-regulated genes), FOS, FOSB and FOSL1, JUNB and JUND (P-Value=3.91E-03, 16% the promoter regions of NGF up-regulated genes; **Fig. 1E**), all belonging to the Reactome biological pathway of NGF-stimulated transcription (HSA-9031628). Fewer motifs were enriched within the 1,000 nucleotide promoter regions of transcripts up-regulated in neurons whose axons were exposed to NT3, and they all related to embryonic development. Such motifs were bound by SOX2, POU5F1 (2.26E-02, 8% of the promoter regions of NT3 up-regulated genes) and SOX17 (P-Value=2.39E-02, 23% of the promoter regions of NT3 up-regulated genes; **Fig. 1F**).

Alternative Polyadenylation (APA) takes place principally co-transcriptionally, giving rise to transcripts expressing identical coding regions and 3’UTRs of often widely different length [21]. To ask whether neurotrophins induce distinct co-transcriptional APA regulation, we investigated the shifts of the relative 3’ UTR usage of transcripts expressed in cell bodies of neurons whose axons were exposed to either NGF or NT3. 27 transcripts showed distal-to-proximal promoter 3’ UTR shift with NGF compared to NT3 (**Supplementary Table 4**), and 45 distal-to-proximal promoter 3’ UTR shifts were observed in response to NT3 (**Fig. 1G**, **Supplementary Fig. 1C** and **Supplementary Table 5**). Tracks of representative transcripts expressing long or short 3’ UTR in cell bodies of neurons treated with NGF or NT3 are shown in **Supplementary Fig. 1D.** Differential 3’ UTR isoform usage can be due to selective increase of promoter-proximal 3’ UTR isoforms, promoter-distal isoforms or both (**Fig. 1H**). Neurons exposed to NGF exhibited a similar number of promoter-proximal and promoter-distal 3’ UTR shifts (20/26). In contrast, NT3 induced twice as many promoter-proximal shifts as promoter-distal shifts (34/16), indicating a preferential usage of isoforms with short 3’ UTR in NT3-treated conditions (**Fig. 1G**). Distal-to-proximal promoter 3’ UTR usage is often indicative of higher translational efficiency [22]. We found that in sympathetic neurons, 3’UTR shift targeted genes associated with distinct biological pathways. Whereas cellular stress and DNA damage were overrepresented terms in response to NGF (**Fig. 1I***, upper panel*), in neurons treated with NT3 target genes were enriched in cell migration and differentiation biological pathways (**Fig. 1**, *lower panel*). Together, these data indicate that NGF and NT3 applied to distal axons are equally capable of regulating gene expression and co-transcriptional APA. This result was expected for NGF, given that NGF/TrkA complexes are retrogradely transported to the nucleus to activate transcription [23–25]. However, it was surprising to see that NT3 was equally capable of eliciting robust transcriptional changes, as NT3 cannot be endocytosed within signalling endosomes and is considered to trigger mostly local signalling [6]. Together, our results demonstrate that NT3 is as effective as NGF in signalling from the axon terminals to the nucleus and is an equally potent regulator of gene expression.

### Identification of RBPs regulating transcript APA in response to NGF and NT3

RNA Binding Proteins (RBPs) have important functions in regulating mRNA processing and metabolism, including alternative polyadenylation [14, 26]. To identify regulators of neurotrophin-specific 3’ UTR APA we interrogated publicly available cross-linking and immunoprecipitation (CLIP) sequencing data for 126 RBPs assayed in human cell lines (**Supplementary Table 6**). Visual inspection of the distribution of RBP cross-link events along the 3’ UTR isoforms identified by our screen revealed that a 500 nucleotide (nt) region preceding the 3’ ends exhibited the highest percentage of bound RBPs (70%) (**Supplementary Fig. 2A**). This region is expected to serve a regulatory function given its high conservation score (**Supplementary Fig. 2B**).

To identify global regulators of APA we searched for associations between cross-linking events mapping to defined regions along the promoter-proximal and promoter-distal 3’ UTRs, and the relative usage in cell bodies of the short 3’ UTR isoform. 126 RBPs were tested individually both in NGF and NT3 conditions (see Material and Methods for details). We found that a positive association between RBP binding and the expression of 3’ UTR isoforms predominantly occurred at the 3’ end of both the promoter-proximal (Ip) and the promoter-distal 3’ UTR (Id), irrespective of the neurotrophin used (**Fig. 2A**). We identified 17 RBPs, preferentially bound within the [-350:+150] region surrounding the 3’ end, as positive regulators of polyadenylation (**Supplementary Fig. 2C,D** and **Supplementary Table 7**). They included the cleavage and polyadenylation (CPA) factors CPSF160 and CSTF2, that are known to reside in the [-50:0] and [0:50] nt regions around the 3’ end [27] (**Fig. 2B,C**). Notably, the binding of RBPs to specific 3’ end terminal regions was positively associated to both short and long 3’ UTR isoforms, which is consistent with the fact that these factors may promote 3’ end processing irrespective of 3’UTR length [28].

**Figure 2.**
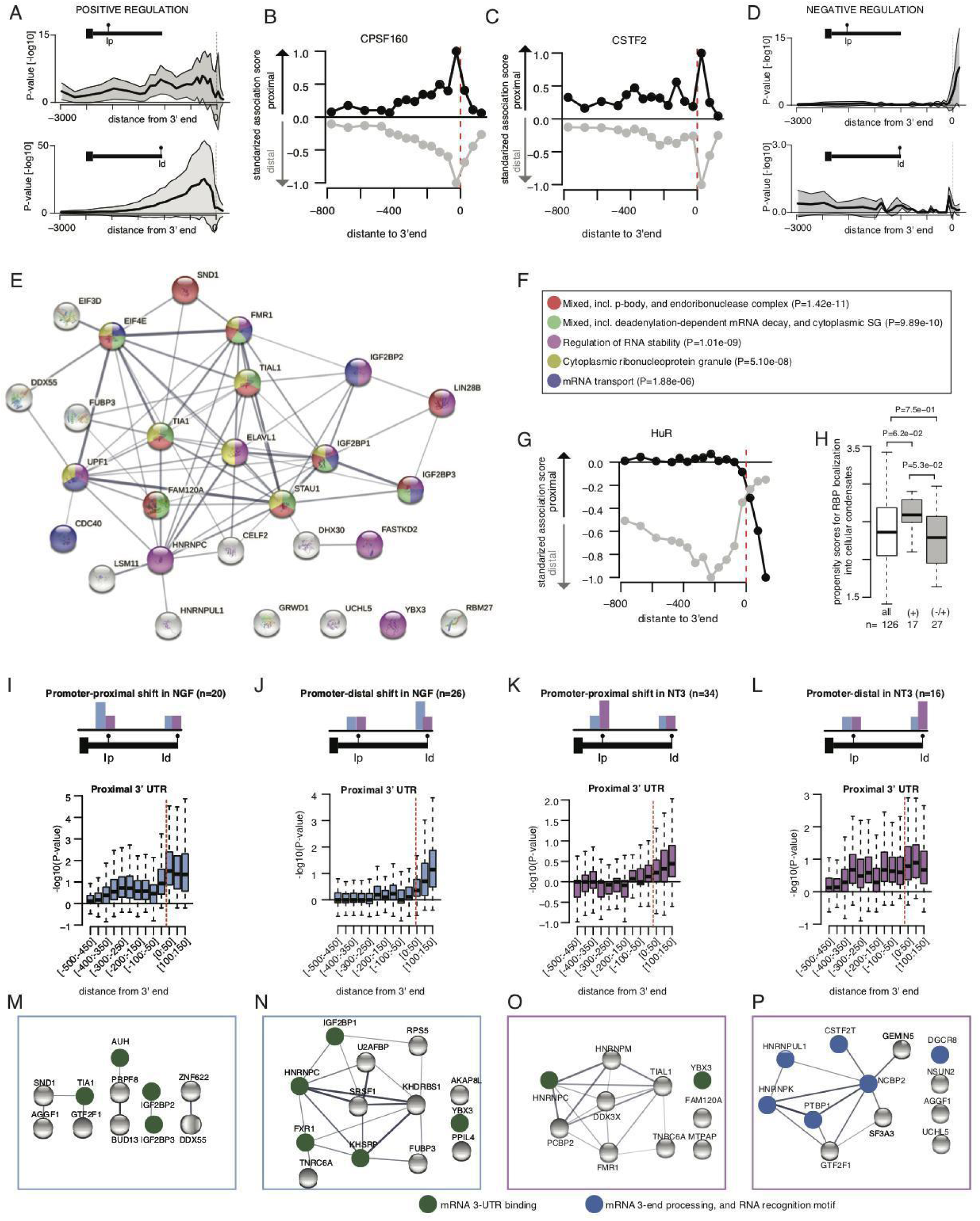
RBP regulome underlying alternative polyadenylation. **(A)** Distribution of significant positive association between the binding of 126 RBPs in defined regions along the 3’ UTR and the relative usage of the promoter-proximal (*upper*) or promoter-distal (*lower*) 3’ UTR isoform. Dark lines display the median significance and shaded areas indicate lower and upper quartiles. **(B, C)** Scatterplot of the extent of standardized significant association between CPSF160 (C) or CSTF2 (D) cross-link event in defined regions along the 3’ UTR and the relative usage of the promoter-proximal (black) or promoter-distal (grey) 3’ UTR isoform in the axons. **(D)** Negative association between RBP binding and the relative usage of the promoter-proximal (*upper*) or promoter-distal (*lower*) 3’ UTR isoform. **(E)** Network of protein-protein interactions for 27 candidate regulators of APA predicted to prevent promoter-proximal usage when bound to the [0:150] nucleotide region down-stream the 3’ end. Edges represent experimentally determined protein-protein interactions annotated in the STRING database ^63^. Nodes indicate proteins coloured according to biological pathways they are enriched in. (**F**) Top five biological pathways and associated P-values over-represented in the 27 candidate negative regulators of short 3’ UTR isoform (Fisher enrichment test). **(G)** Same as (C) for HuR. **(H)** Distributions of the propensity scores of the 126 RBPs, the 17 positive regulators of APA (short and long 3’ UTR isoform) and the 27 negative regulators of the short 3’ UTR isoform to localize into cellular condensates as predicted by GraPES . (Welch’s t-test assessing the significant difference between the mean propensity scores). (**I-L**) Distribution of Fisher enrichment scores (-log10(P-value)) in RBP binding along the short 3’ UTR isoforms of the pairs of isoforms exhibiting significant promoter-proximal shift () or promoter-distal shift (I and J) in NGF, and promoter-proximal shift or promoter-distal shift (K and L) in NT3. (**M-P**) Network of protein-protein interactions for RBPs exhibiting significant enrichment in cross-link events in the [0:+150] nt regions downstream the 3’ end of the short 3’ UTR isoforms associated with significant promoter-proximal shift (M) or promoter-distal shift (N) in NGF and promoter-proximal shift or promoter-distal shift (O and P) in NT3. Edges represent experimentally determined protein-protein interactions annotated in the STRING database. Nodes indicate proteins coloured according to biological pathways they are enriched in.

Negative association between 3’ UTR isoform usage and RBP binding revealed that negative regulators of polyadenylation are uniquely detected in the [0:150] nt region downstream of the promoter-proximal 3’ end. Long 3’ UTR isoforms were not associated with any negative regulator (**Fig. 2D**). Interestingly, the 27 negative regulators of the short 3’ UTR form a densely connected network of experimentally validated interacting proteins that are enriched in biological processes related to mRNA stability (**Fig. 2E,F, Supplementary Fig. 2E,F** and **Supplementary Table 8**). HuR is a well characterised RBP that competes with the CSTF factors downstream of poly(A) sites to block polyadenylation [29]. Accordingly, we found that HuR binding downstream of the 3’ end of the promoter-proximal 3’ UTRs strongly associated with decreased promoter-proximal 3’ UTR usage, promoting the transcription of long 3’ UTR isoforms. In contrast, HuR binding upstream of the 3’ end does not affect polyadenylation (**Fig. 2G**). Thus, the choice of short or long 3’ UTR mainly relies on factors residing within the 150 nt region downstream of the promoter-proximal 3’ end. Analysis of the predicted propensity to localise into cellular condensates revealed significant differences between the 17 positive and the 27 negative regulators of polyadenylation, with the former exhibiting higher probability to localise into P-bodies and cytoplasmic granules (**Fig. 2H**). Altogether this analysis identifies novel and previously known regulators of polyadenylation along with their preferential location around the 3’ end. The data also show that positive regulators are more likely to concentrate into condensates where they may operate to promote the cleavage of the 3’ end [10].

Next, we asked whether NGF and NT3 may utilise distinct regulators of 3’ UTR APA. Although exposure of distal axons to NGF or NT3 resulted in very similar APA profiles in the cell body, subtle but significant differences were observed in dozens of transcripts (**Fig. 1F**). 3’ UTR position-dependent Fisher enrichment in RBP cross-link events were studied for the four groups of APA isoforms previously identified (**Figs. 1F,G**). The distributions of the enrichment scores (-log10(P-value)) along the 3’ UTR revealed that the regulatory regions with significant enrichment of RBPs were restricted to the region downstream the promoter-proximal 3’ end, irrespective of the grouping (**Figs. 2I-L** and **Supplementary Fig. 2G**). This result suggests that the selection of either isoform between NGF and NT3 occurs through the binding of RBP factors that compete with cleavage factors in the downstream region of the 3’end. This mechanism is similar to the one predicted to promote the long 3’ UTR in lieu of the short. Significantly enriched regulators (P-value <0.01) included four networks of RBPs associated with mRNA 3’ UTR binding and 3’ end processing (**Fig. 2M-P**). Altogether our data identify neurotrophin-specific regulators of 3’ UTR APA in sympathetic neurons by integrating human-derived CLIP data with rat sequencing data.

### NGF and NT3 induce the localisation of distinct mRNA transcripts in axons

We next asked whether NGF and NT3 induced the localisation of distinct mRNAs in axons. A similar number of transcripts and 3’ UTR isoforms were found in cell bodies and axons independently of neurotrophin treatment (**Fig. 3A** and **Supplementary Fig. 3A**). However, a significant number of axonal mRNAs with distinct cellular functions (**Supplementary Fig. 3B**) were uniquely detected in response to either NGF (n=1,962) or NT3 (n=1,089; **Fig. 3B**). In line with previous studies [10, 30], a larger percentage of axonal transcripts expressed multiple (**Fig. 3C**) and longer (**Fig. 3D**) 3’UTR isoforms, when compared to cell bodies in both conditions.

**Figure 3.**
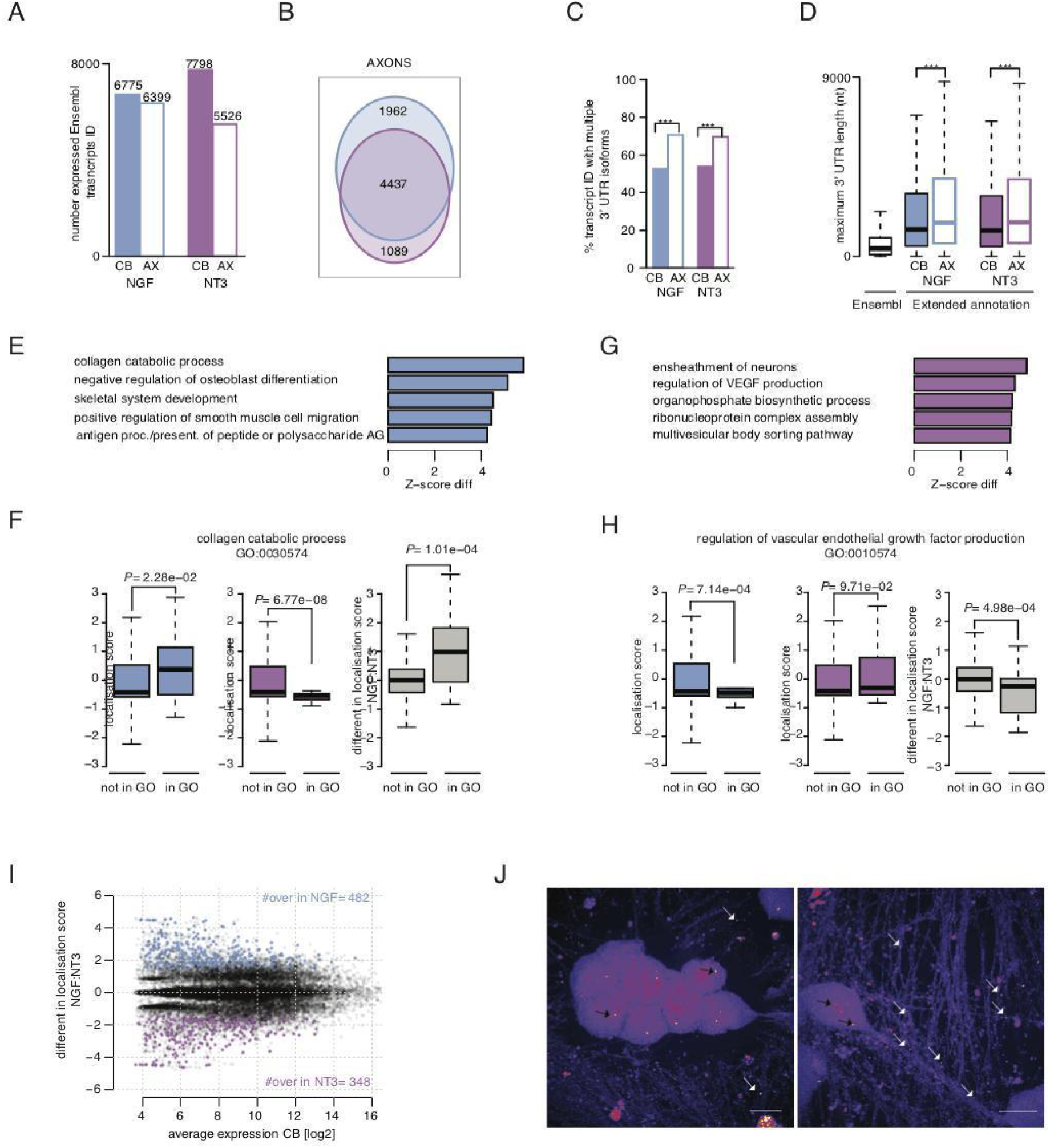
Sympathetic neuron axons exposed to NGF or NT3 contain distinct 3’ UTR isoforms. **(A)** Ensembl transcripts ID detected in cell bodies only (full bars), and cell bodies and distal axons (empty bars) in NGF (blue) and NT3 (purple) culture conditions. **(B)** Overlap between the transcripts detected in the distal axons of neurons exposed to either NGF or NT3. **(C)** Cell body (full bars) and axonal (empty bars) transcript IDs showing multiple 3′ UTRs in either NGF or NT3 culture conditions. (***p <0.001; two-sided Fisher’s exact count test). **(D)** Maximum 3′ UTR lengths for existing annotations in Ensembl Rn5 (Ensembl) and for those newly identified by 3′ end RNA sequencing (Extended annotation) in either NGF or NT3. (***P<0.01; two-sided Wilcoxon rank-sum test). **(E)** Top five GO biological pathways which associated transcripts exhibit the most significant increase in axonal localisation in NGF (*left)* compared to NT3 (*right)*, as quantified with standardized scores comparing GO-annotated transcript with the full pool of detected transcripts (Z-score). **(F)** Distributions for the background genes and the genes belonging to the collagen catabolic biological pathway of the localisation scores in NGF (*right*), the localisation scores NT3 (*center*), and the differences in localisation scores between the two conditions (*right*). (Welch’s t-test assessing the significant difference between the mean localisation scores. Boxplots display the five number summary of median, lower and upper quartiles, minimum and maximum values). **(G)** Same as (F) for the top five GO biological pathways exhibiting excess axonal localisation in NT3 compared to NGF. **(H)** Same as (G) for genes belonging to the regulation of VEGF production. **(I)** Average mRNA abundance across the four cell bodies and the difference in localisation scores between NGF and NT3. 482 transcripts with significantly more localized in NGF as compared to NT3 (blue dots; Z-score>1.96) and 348 transcripts significantly more localized in NT3 compared to NGF (purple dots; Z-score<-1.96). **(J)** Lower levels of Atf3 localized to NGF-treated axons (*left*) compared to NT-3 treated axons (*right*). Black arrows point to cell body signal, white arrows point to axonal signal. Scale bar, 10μm.

The abundance of axonal mRNAs mostly correlated with expression levels in the cell bodies and transcript length (**Supplementary Fig. 3C,D**), suggesting that diffusion and anchoring are important mechanisms regulating mRNA localisation in axons. To identify the regulatory mechanisms responsible for active mRNA axonal localisation, we developed a statistical model and a novel metric that we named Localisation Score (LS). LS quantifies the efficiency of transcripts localisation in axons irrespective of mRNA length and abundance in the cell bodies (**Supplementary Fig. 3E**; **Supplementary Table 9**; see Material and Methods). Thus, LS provides a better statistical tool than the gene abundance ratio between axons and cell bodies commonly used to quantify mRNA localisation, and enables the identification of over-transported mRNA irrespective of the expression levels (**Supplementary Fig. 3F,G**). A positive LS indicates higher axonal mRNA abundance than would be expected for transcripts with similar size and expression level and correlates with active transport and stabilisation. Conversely, negative LS values are indicative of either transcript diffusion from the cell bodies or higher mRNA degradation. Using this novel metric, we identified thousands of over- and under-transported 3’ UTR isoforms in both NGF and NT3 conditions (**Supplementary Fig. 3H-J**). To validate the statistical model, real-time quantitative PCR (RT-qPCR) was performed on transcripts predicted to be restricted to cell bodies. Analysis of Eid2 and Rab22a indicated that the mRNAs were virtually absent in NGF and NT3 treated axons, despite being highly expressed in the cell bodies (**Supplementary Fig. 3K,L**). LS analysis revealed that NGF and NT3 promote axonal localisation of transcripts associated with largely similar GO biological pathways, such as vesicular localisation and axo-dendritic transport (**Supplementary Fig. 3M,N**). Subtle but significant differences in axonal localisation of transcripts associated to specific biological pathways were detected in both conditions. For example, transcripts related to the collagen catabolic pathway exhibited higher LS in NGF axons compared to NT3 (**Fig. 3E,F**), while those related to the vascular endothelial-derived growth factors-related pathway had higher LS in NT3 (**Fig. 3G,H**). The latter finding is especially interesting considering that during the early development when sympathetic neurons are exposed to NT3, axons grow in close contact with blood vessels [7], possibly inducing the transport of transcripts that mediate cross-signalling between neurons and endothelial cells. Analysis performed on individual transcripts identified 482 3’ UTR isoforms with significantly higher LS in NGF axons and 348 with higher LS in NT3-treated axons (**Fig. 3I** and **Supplementary Tables 10,11**). Single molecule RNA Fluorescence In Situ Hybridization (smFISH) of *Atf3*, a transcript predicted to localise in NT3 but not NGF-treated axons, confirmed that neurotrophins induce mRNA axonal localisation in a highly specific manner (**Fig. 3J**) [31]. These findings identify a large repertoire of transcripts that are localised and stored in developing sympathetic neuron axons. They also indicate that both NGF and NT3 applied to distal axons select the transcripts that will be localised peripherally.

### Comprehensive identification of the RBP regulome for axonal mRNA localisation

The interaction of *cis*-regulatory elements with *trans*-acting factors such as RBPs plays a crucial role in regulating mRNA stability and axonal transport [2, 32]. We next investigated whether specific RBPs accounted for the distinct mRNA axonal localisation observed in NGF and NT3 treated axons. Using the CLIP data described above, LS of transcripts with cross-linking events was tested for each individual RBPs on regions of 50 nucleotides along the 3’ UTR (see Material and Method). We found that the [-200:-50] nt region preceding the 3’ end exhibited the highest regulatory potential for axonal localisation in both NGF and NT3 conditions (median P-value=10e-25 across the 126 RBPs; **Fig. 4A**). The preferential binding of axonal localisation *trans*-acting factors to this specific region is in line with previous findings demonstrating the presence of NGF-dependent localisation elements within -150 nt from the 3’ end [33]. Indeed the number of RBPs with cross-link events within the [-150:-100] nt region preceding the 3’ end correlated with the LS for the isoform (**Fig. 4B**). The 32 RBPs exhibiting the highest positive regulatory association with localised transcripts regulate mRNA transport (**Fig. 4C** and **Table S12**). Experimentally validated regulators of transport, such as FMR1 (**Figs. 4D**), IGF2BP1 [33] (**Supplementary Fig. 4A**) and TDP43 [34, 35] (**Supplementary Fig. 4B**), exhibited significantly higher cross-linking events in over-transported, compared to under-transported transcripts in both NGF and NT3 conditions. Although 126 RBPs showed similar positive association with axonal localisation in NGF and NT3 axons, (**Supplementary Fig. 4C**), SmB (**Fig. 4E**) and EIF4G2 (**Supplementary Fig. 4D**) showed significant enrichment in cross-linking events in over-transported 3’ UTR isoforms in either NGF or NT3 conditions, respectively. eIF4G2 is especially relevant given that it is locally translated in rat sympathetic neuron axons to support axon growth [36].

**Figure 4.**
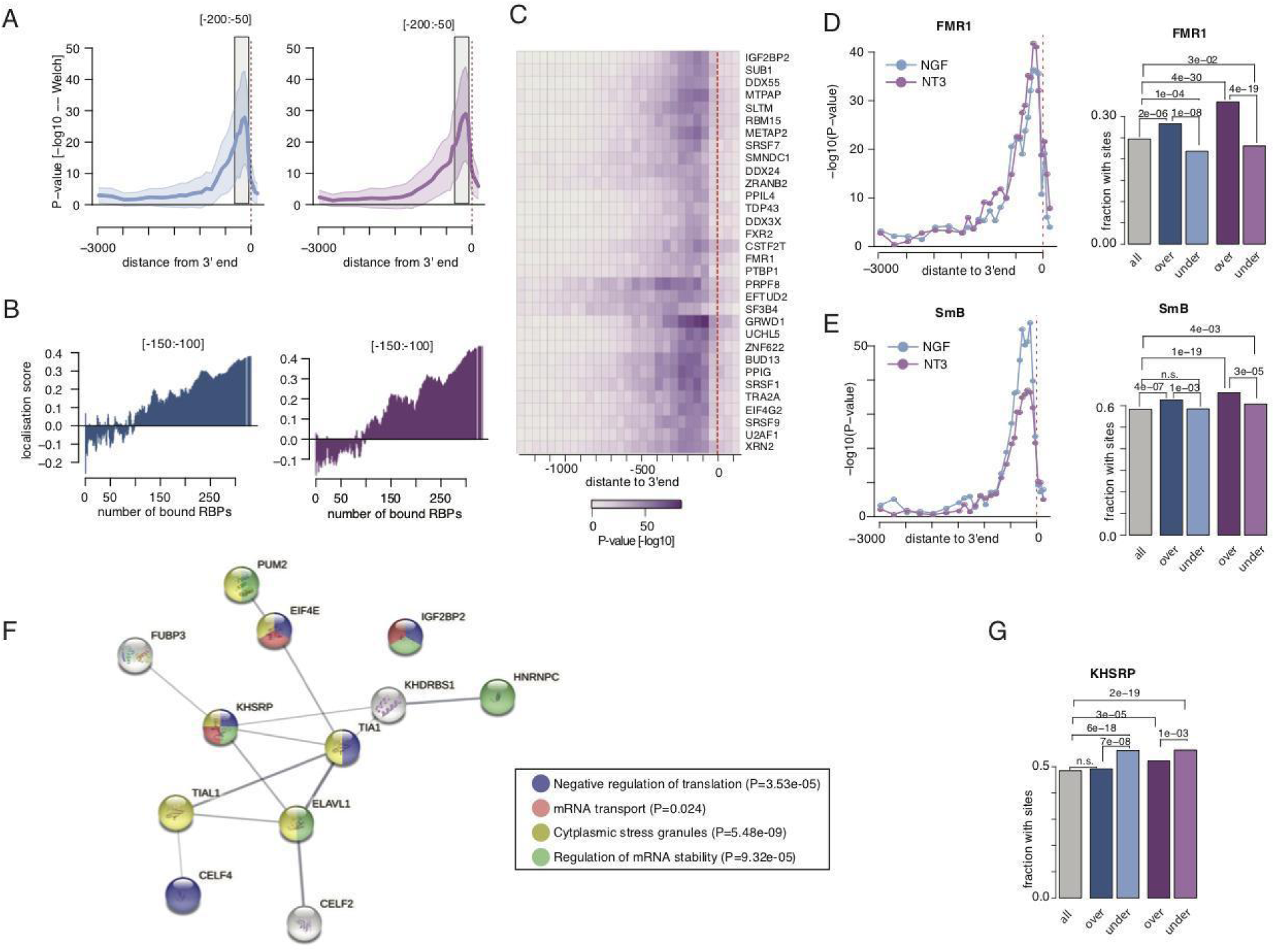
The RBP regulome underlies the axonal localisation in NGF and NT3. **(A)** Distribution of the significance in the difference between the mean localisation scores in NGF (*left*) and NT3 (*right*) of the isoforms exhibiting or not a cross-link event in 50 nucleotide long regions along the 3’ UTR for 126 RBPs. Dark line displays the median and shaded areas indicate lower and upper quartiles. **(B)** Increasing localisation scores in NGF (*left*) and NT3 (*right*) conditions as a function of the number of RBPs with cross-link events in the [-150:-100] nucleotides upstream the 3’ end. **(C)** Heatmap showing the significance in the difference between the mean localisation scores of the 3’ UTR isoforms exhibiting or not a cross-link event in 50 nucleotide long regions along the 3’ UTR for the top 32 RBPs positively associated with axonal localisation. **(D)** Scatterplot of the extent of significant association between FMR1 cross-link event and the axonal localisation along the 3’ UTR (*left)*. Blue = NGF; purple = NT3. (*Right*) Barplots showing the fraction of [-250:-50] nucleotide regions upstream the 3’ ends exhibiting FMR1 cross-link event in the full set of detected transcript (grey bar), and the pools of over- and under-transported transcripts in NGF (blue bars) and NT3 (purple bars), respectively. (Fisher enrichment test). **(E)** Same as (E) for SmB. **(F)** Protein-protein interactions for 16 RBPs exhibiting significant enrichment in cross-link events in the [-250:-50] nt regions upstream the 3’ end of the under-transported transcripts. Edges represent experimentally determined protein-protein interactions annotated in the STRING database. Nodes indicate proteins coloured according to biological pathways they are enriched in. Inset: top 4 biological pathways and associated enrichment P-values as obtained from Fisher enrichment test. **(G)** Same as (E) for KHSRP.

Although negative RBP regulator of axonal localisation were not found using the Welch’s t-test comparing the LS between RBP targets and others, 12 RBPs showed significantly higher binding occurrence (Fisher count test) in the [-250:-50] nucleotide region preceding the 3’ end of the under-transported isoforms, compared to either all transcripts or over-transported isoforms (**Supplementary Table 13**). These RBPs form a connected network of interacting proteins enriched in regulators of translation and mRNA stability (**Fig. 4F**), including HuR/Elavl1 that decreases neuronal cytoplasmic mRNA, CELF4 and KHSRP [37–39] (**Fig. 4G** and **Supplementary Fig. 4E,F)**. Both KHSRP and CELF4 regulate mRNA abundance in axons and dendrites, and KHSRP localises to both axons and dendrites [40]. This finding supports the hypothesis that mRNA destabilisation may play a key role in regulating the expression of localised transcripts [39, 41]. Notably, we did not find RBPs linked to decreased localisation of specific axonal mRNAs in response either to NGF or NT3. Taken together, these findings indicate that mRNA stability is likely to be a key component underlying limited axonal localisation.

### The RBPome for 3’UTR remodelling in axons

We recently discovered that the 3’UTR of transcripts localised in axons may undergo local remodelling, resulting in the generation of a polyadenylated isoforms expressing a shorter 3’UTR that are efficiently translated, and stable, 3’UTR-derived fragments with yet unknown functions [10, 20]. Analysis of isoforms localised in NGF and NT3-treated axons revealed a similar number of 3’UTR isoforms with different usage of long and short 3’UTRs in both cell bodies and axons (665 and 458, respectively; **Supplementary Figs. 5A,B; Supplementary Table 14-17**). Candidate transcripts of axonal remodelling were identified in both NGF and NT3 conditions as isoforms expressing short 3’UTR abundant in axons but virtually absent in cell bodies (**Fig. 5A**, *left*; **Supplementary Table 18-19**). This unique expression pattern suggests that they are not the product of co-transcriptional APA and are cleaved in axons. Interestingly, isoforms that express short 3’UTRs only in axons were largely not overlapping between NGF and NT3 (**Fig. 5A**, *right*), suggesting substantial differential regulation between these two conditions. Tracks of representative transcripts expressing unique short 3’ UTR in axons treated with either NGF or NT3 are shown in **Fig. 5B,C**.

**Figure 5.**
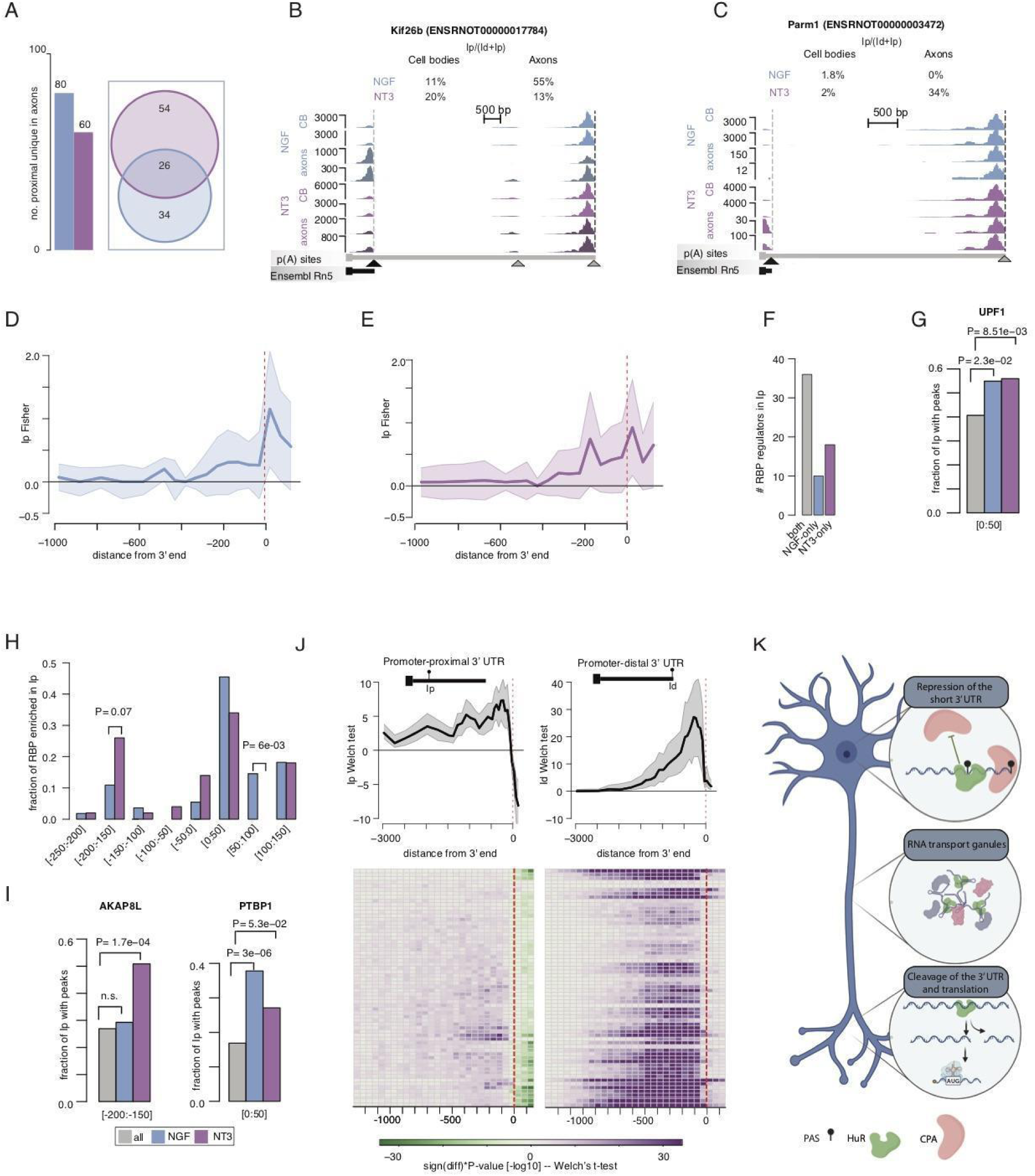
Negative regulators of APA in the cell body are candidate 3’ UTR cleavage factors in the axons. **(A)** Number of 3’UTR isoforms with proximal shift uniquely detected in axons i.e. candidate to remodeling in NGF (blue) and NT3 (purple) culture condition (*left*). Venn diagram showing the overlap between the candidate of axonal remodeling in NGF and NT3 (*right*). **(B)** A representative transcript with a marked shift toward axonal increase in promoter-proximal 3’ UTR uniquely detected in axons treated with NGF (Kif26b). **(C)** A representative transcript with a marked shift toward axonal increase in promoter-proximal 3’ UTR uniquely detected in axons treated with NT3 (Parm1). **(D, E)** Distribution of the extent of significant enrichment in 126 RBP cross-link events in defined regions along the 3’ UTR of the 80 (**D**) and 60 (**E**) candidate isoforms of axonal remodeling in NGF and NT3 condition, respectively. Dark lines display the median significance and shaded areas indicate lower and upper quartiles. **(F)** Number of candidate RBPs regulators of axonal remodeling identified in the [-250:+150] nt region surrounding the 3’ end of the promoter-proximal isoform in both conditions (gray bar), in NGF only (blue bar) and NT3 only (purple bar). **(G)** Fraction of promoter-proximal 3’ UTR isoforms exhibiting cross-link events for UPF1 in [0:50] nucleotide region down-stream of the cleavage site. Gray bar = all promoter-proximal 3’ UTR; blue bars = 80 candidates of axonal remodeling in NGF; purple bars = 60 candidates of axonal remodeling in NT3. **(H)** Fraction of candidate RBPs regulators of axonal remodeling in NGF and NT3 at specific positions along the 3’ UTR where they exhibit the most significant enrichment. (Fisher enrichment test). (**I**) Same as (G) for AKAPL and PTBP1. **(J)** Distribution of the extent of significant positive or negative association between the cross-link events in defined regions along the short (*upper left*) and long (*upper right*) 3’ UTR isoforms of 65 candidate regulators for axonal remodeling, and their relative usage of the short and the long isoform in the cell body. Dark lines display the median significance and shaded areas indicate lower and upper quartiles. Heatmaps showing the individual significance association between cross-link events of individual RBPs along the short (*lower left*) or long (*lower right*) 3’ UTR isoform and their relative usage in the axons. **(K)** Proposed model where the local release of mRNAs and cleavage factors from transport granules may serve as a macromolecular mechanism linking co-transcriptional APA, RNA localization, axonal 3’ UTR remodeling to translation and neurotrophic-mediated axon growth.

We recently identified Argonaute 2 (Ago2) as the endonuclease responsible for 3’UTR cleavage [10], however how 3’UTR remodelling activity is regulated in axons remains unknown. Fisher enrichment in RBP cross-link events along the 3’ UTR of the remodelled isoforms revealed that while NGF-related predicted regulators of axonal remodelling preferentially localised to the [0:+50] nt region downstream the 3’ end, NT3-related predicted regulators were found on both sides of the 3’ end (**Figs. 5D,E**). Thirty-five enriched RBPs were observed in both NGF and NT3 remodelled isoforms, whereas 10 and 18 were unique for NGF or NT3, respectively (**Fig. 5F** and **Supplementary Table 20-21**). Common RBPs included HuR which is known to inhibit polyadenylation when bound to the region downstream of the 3’ end (**Supplementary Fig. 5C**). UPF1, which belongs to the protein complex that we previously showed to mediate the 3′ UTR cleavage [1], was significantly enriched in the 50 nt region downstream of the 3’ end of remodelled isoforms in both NGF and NT3 treated axons (**Fig. 5G** and **Supplementary Fig. 5D**). Analysis of the positional preference of the top predicted regulators of axonal cleavage revealed that for NT3-remodelled isoforms binding to the [-200:-150] nt window upstream the 3’ end is favoured, whereas regulatory regions of NGF-remodelled isoforms reside in the [50:100] nt region downstream the 3’ end (**Fig. 5H**). Neurotrophin-specific predicted regulators of axonal remodelling include AKAP8L, which is enriched in the [-200:-150] nt region of 3’ UTR short isoforms and is uniquely detected in NT3 condition, and PTBP1, which is enriched in the [0:50] nt region downstream of the pool of short 3’ UTR isoforms detected only in NGF-treated axons (**Fig. 5I**).

We next studied whether RBPs predicted to regulate 3’ cleavage in axons may dictate the choice of the 3’ end in the nucleus. The analysis revealed that these factors when bound downstream of the 3’ end of short 3’ UTR isoforms, behaved as negative regulators of short 3’ UTR isoforms (**Fig. 5J**). Based on these findings, we propose a model in which RBPs responsible for axonal cleavage are primarily recruited to the downstream region of the promoter-proximal 3’ end in the nucleus, where they favour the expression of the long 3’ UTRs required for axonal localisation possibly by competing with cleavage factors. Once they reach the distal part of the axons, they promote the cleavage of the long 3’UTR by probably interacting with additional RBPs such as the ones we previously identified [1] (**Fig. 5K**).

## Discussion

### NT3 and NGF applied to distal axons regulates transcription

Despite binding to the same receptor TrkA, NGF and NT3 intracellular signalling is quite distinct. NGF-TrkA complexes are internalised into signalling endosomes that travel back long distances to the cell bodies where they activate transcription. In contrast, NT3 is a growth factor released by target tissues mostly during the early stage of neuronal development when axons are still short, and it is thought to act mostly locally in axon growth cones [6–9, 42]. Our data demonstrate that both neurotrophins when applied to distal axons, induce a robust transcriptional response (**Fig.1**). We also found that NGF and NT3 applied to distal axons regulate APA, inducing the expression of 3’UTR isoforms that are only partially overlapping. Thus, our findings indicate that NT3 is as capable as NGF of inducing transcriptional changes, opening the possibility that despite not being internalised in signalling endosomes, NT3 initiates a signalling pathway that is faithfully propagated to the nucleus. Given that the modality of how NGF containing signalling endosomes communicate to the nucleus is unknown, it is conceivable that both neurotrophins may share common intranuclear regulatory pathways, perhaps involving changes of nuclear calcium concentration [43].

### The RBPome for axonal localisation

Subcellular localisation of mRNA enables the synthesis of proteins necessary for axon growth and synaptic functions [2, 44, 45]. Here, we show that axonal exposure to NGF and NT3 regulates axonal localisation of specific transcripts enriched in defined biological pathways (**Fig. 3**). RBPs are key regulators of mRNA metabolism including mRNA transport and translation, and they are essential in determining when and where specific proteins will be expressed. By integrating our 3’ end-seq data with CLIP data generated in human cell lines, we discovered novel and known regulators of APA (**Fig. 2**), corroborating the suitability of human data to study the RBP regulome underlying mRNA metabolism in rat neurons. Using these data, we found that axonal localization is positively regulated by RBPs that facilitate mRNA transport, and negatively modulated by RBPs that regulate mRNA stability. Albeit universal axon-targeting motifs have not been identified so far, 32 RBPs were found to positively associate with axonal localisation (**Fig. 4**). Furthermore, only minor differences in predicted regulators of transport were observed between NGF and NT3 treated axons, and no evidence was found for differential binding of destabilizing proteins and possibly mRNA stability between the two conditions. Thus, although NGF and NT3 regulate the transport of specific mRNA isoforms to axons, they utilise similar RBP complexes, suggesting a conserved mechanism for transcript localisation in neurons. Future studies will clarify whether the same combination of RBPs drives the transcript localisation to other neuronal compartments, such as dendrites and dendritic spines.

### A switch of RBP complex composition promotes 3’ UTR remodelling in axons

We recently discovered that few axonal transcripts including *Inositol Monophosphatase 1* (*IMPA1*) undergo 3’UTR remodelling in sympathetic neuron axons [20]. Importantly 3’UTR cleavage is necessary for triggering the translation of *IMPA1* mRNA isoforms expressing a short, cleaved 3’UTR [20]. Hundreds of transcripts with a proximal-to-distal shift of the 3’UTR are detected in axons. Several short 3’UTR isoforms were expressed exclusively in axons and in some cases specifically in response to either NGF and NT3 (**Fig.5** and **Supplementary** **Fig.5**). Analysis of the RBP complexes interacting with the remodelled isoforms revealed enrichment of UPF1 and PTBP1 that we previously identified as binding partners of the 3’UTR remodelling complex [20]. Based on our data, we propose that a group of RBPs is recruited co-transcriptionally to the region downstream of the proximal 3’ UTR competing with CSTF factors and suppressing the use of the proximal PAS. This event results in 3’ UTR lengthening, which is compatible with several studies indicating that isoforms expressing long 3’UTR are preferentially transported to axons [15, 20, 46]. These factors may remain bound to the 3’ UTR within RBP granules and hitchhike along the axons to eventually co-localise in the axonal compartment with the long 3’ UTR isoform. Upon de-assembly of the transport granules in axons, some factors, including UPF1, promote the cleavage of the long 3’UTR into the short 3’UTR isoform. Transport granules have been considered as “translation factories” that contain RBPs, mRNAs, ribosomes and translation factors, and regulate local protein synthesis [47, 48]. Here, we propose a similar mechanism by which APA, RNA localization and neurotrophin-dependent translation are coupled and co-regulated. In axons, release of 3’ UTR isoforms remodelling factors from granules may serve as the final step that allows translational activation of proteins expressing a shorter 3’UTR necessary for axon growth (**Fig. 6F**).

## CONCLUSIONS

Our study shines new light on the nature of both the mRNA localised in axons and the RBPs that regulate their localisation and 3’UTR remodelling. Many neurological diseases are now considered as disorders of the RNA [19, 49]. Our data provide a wealth of potentially new druggable targets that may open novel therapeutic avenues for the cure of degenerative and traumatic disorders of the central and peripheral nervous system.

## MATERIALS AND METHODS

### Reagents

Cell culture reagents, molecular biology reagents and kits were purchased from Thermo Fisher Scientific and all other chemicals from Sigma, unless stated otherwise.

### Compartmentalized cultures of rat sympathetic neurons

All animal studies were approved by the Institutional Animal Care and Use Committees at University College London. Superior cervical ganglia were dissected from post-natal day 1 (P1) Sprague Dawley rats, enzymatically dissociated and plated on glass coverslips or in compartmentalized chambers pre-coated with home-made collagen and laminin (5μg/mL), as previously described [23]. When using compartmentalized chambers, NGF was reduced to 10ng/mL in the central compartment five to six days after plating and neurons were then maintained with either NGF (100 ng/ml) or NT3 (1μg/mL) in the lateral compartment. Cytosine arabinoside (ARA-C, 10μM) was added 24 hours after plating to block the proliferation of non-neuronal cells.

### RNA isolation, reverse transcription, linear amplification and 3’end-RNASeq

To ensure that the axons were free of cell bodies, prior to each experiment axon compartments were incubated with Hoechst 33342 (10μg/mL in PBS for 20 min at 37°C) and observed under an inverted fluorescent microscope. Cultures showing cell nuclei in the axon compartments or leakage of the dye in the central compartment were discarded. Total axonal and cell bodies RNA was purified from the lateral compartments of 52 or 36 chambers and the central compartment of 7 or 6 chambers respectively obtained from 3 or more independent cultures. Total RNA was isolated using PureLink® RNA Micro Scale Kit, according to the manufacturer’s instructions with minor modifications. Briefly, axons and cell bodies were collected from chambers using a lysis buffer (300μL) containing 10%β-mercaptoethanol. Total mRNA bound to the columns was washed and eluted twice in elution buffer (12μL). Aliquots of each sample were reverse transcribed in a 20μL reaction volume containing random hexamer mix and 50U SuperScript III Reverse Transcriptase at 50°C for 1 hr. To check the quality of samples and the absence of cell bodies contamination in axon samples, first-strand cDNAs (5μL) were PCR amplified in a 25μL PCR reaction containing actin beta or histone H4 specific primers (0.20μM), dNTPs (200 nM) and Go Taq polymerase (1.25U, Promega). Primer sequences and PCR conditions are provided (**Table S23**).

For mRNA linear amplification, samples were purified as described above, concentrated by speed-vacuum centrifugation to 1μL (axons) or 5μL (cell bodies) volume, and used for two rounds of linear amplification as previously described (Baugh et al., 2001). The volume of the first-strand reaction for the axons was scaled down to 5μL. After the second round of amplification contaminant cDNA was digested by treating the samples with RNAse-free DNAse (2U, Epicentre). Performance of the samples was tested by RT-PCR. Linear amplified aRNA from cell bodies and axon samples (2 biological replicates each) was used to prepare RNASeq libraries using the strand-specific ScriptSeq protocol. Paired-end sequencing (2x 150bp) of four indexed libraries was performed on the Illumina HiSeq2000 platform, generating in excess of 80M mappable reads per sample. Library preparation and sequencing were performed at the Liverpool Centre for Genomic Research (CGR, http://www.liv.ac.uk/genomic-research/).

### RT-PCR and quantitative RT-PCR

mRNA was isolated from sympathetic neurons as described above and reverse transcribed. qRT-PCR reactions (25μL) contained 12.5μL of Sybr Select or Luna (NEB) Mastermix and 0.25 μM primers, unless otherwise indicated. Reactions were performed in triplicate on a Biorad CFX Connect Real-time Machine. The Comparative Ct Method (ΔΔCt Method) was used for relative quantification. At the end of 40 cycles of amplification, a dissociation curve was performed in which SybrGreen fluorescence was measured at 1°C intervals between the annealing temperature and 100°C. Peaks of the melting temperatures of amplicons varied between 80°C and 90°C. Primer sequences and PCR conditions are described in **Table S23**.

### Single molecule FISH (smFISH)

Probe sets targeting the 3’UTR of rat *Atf3* (Stellaris probes, Biosearch technologies) were designed using the Stellaris probe-set designer tool to specifically detect the 3’UTR of the transcript and 3’end labelled with CalFluor590. Probes were reconstituted at 12.5μM in TE buffer (10mM Tris pH8, 1mM EDTA pH8). smFISH was performed as previously described [20] on SCG neurons cultured on glass coverslips for 5 days in high NGF concentration (100ng/mL) and then either maintained in high NGF or washed and maintained in NT3 (1μg/mL) for further 7 days before fixation. Cells were washed with PBS, fixed using 3.7% PFA at RT for 10 mins, permeabilised with 70% EtOH at 4°C for 3.5 hrs and then pre-hybridised in 2xSSC 10% Formamide for 5mins at RT. 1μl of 12.5μl probe stock was added to 100μl Hybridisation buffer (10% Dextran Sulfate, 2xSSC, 10% Formamide, 2mM vanadyl ribonucleoside, 0.1ug/uL salmon sperm DNA, 0.1% Triton X-100, 1% BSA) before incubation of the coverslips O/N at 42°C in humidified chamber. Coverslips were then washed 2×30 mins in warm 2xSSC 10%Formamide at 37°C in the dark and 1×5 min in PBS+100ug/mL DAPI before mounting to slides with ProlongGold. Coverslips were cured O/N at RT before imaging at a 3i confocal microscope (Intelligent Imaging Innovations, Inc.) equipped with a Photometrics Prime 95B (Scientific CMOS) camera. Images were processed using ImageJ software and figures prepared using Adobe Photoshop software.

### Inference of 3’UTR isoforms from 3’-end RNA-seq

The rat genome is poorly annotated compared with mouse and human; therefore, 3′ end RNA-seq data were used to identify unknown isoforms by re-annotating the 3′ ends to the Ensembl Rn5 database (v.78) as previously described [20]. Briefly paired-end stranded RNA-seq reads of 150 bp were mapped to the reference rat genome (UCSC, rn5), and nucleotide-level stranded coverage obtained for axons and cell body enabled the identification of continuously transcribed regions. Continuously expressed fragments were next associated with matching strand overlapping 3’UTR using Ensembl version 78 (v78) [50], and extensive downstream filtering was performed to exclude potential intragenic transcription, overlapping transcripts, and retained introns as described in [51]. Segmentor3IsBack R package was finally used to segment the longest 3’ UTR isoform in regions of spatially coherent coverage hence to identify alternative 3’UTR isoforms [52]. 3’ ends located within 50 nt distance were clustered together, selecting the most promoter-distal annotation. The GTF annotation file was deposited together with the source code to generate the new 3’ UTR annotation on GitHub : https://github.com/RLuisier/my3UTRs.

### 3’UTR isoform quantification and identification of transcripts localized to axons

As previously described, the last 500 nt portion of each transcript contains above 70% of the reads originating from that transcript irrespective of their length. We thus used the number of reads mapped to the -500 nt terminal region of each 3’ UTR isoform as a proxy for the 3’ UTR isoform expression levels. The density of mapped reads in -500 nt terminal region of 3’UTR isoforms is bimodal, with a low-density peak probably corresponding to background transcription, i.e. 3’UTR isoforms of low abundance or 3’UTR isoforms to which reads were spuriously mapped, and a high-density peak corresponding to expressed 3’UTR isoforms (**Supplementary Fig. 3A**). In order to identify 3’UTR isoforms expressed in axons and cell body, a two-component Gaussian mixture was fitted to the data using the R package mclust [53]. An isoform was called expressed if there were less than 5% chance of belonging to the background category in both replicates or if there was more than 10% chance of belonging to the expressed category in at least one replicate.

### Differential gene expression analysis

For this analysis we focussed on the data obtained from the cell body compartments. The pipeline described above outputs 3’ UTR transcript abundance, and thus we first calculated the abundance of genes by summing up the estimated raw count of the constituent isoforms to obtain a single value per gene. Differential analysis was performed using the edgeR package [54]. Normalization factors were computed using the TMM technique, after which tagwise dispersions were calculated and subjected to an exact test. Resulting P values were subjected to Benjamini–Hochberg multiple testing correction to derive FDRs. Genes were considered as differentially expressed between NGF and NT3 if log2FC>0.58 (FC>1.5) and P value <0.01. The complete lists of differentially expressed genes are reported in **Supplementary Tables 1 and 2**.

### Transcription Factor Binding Site analysis

The 1000 nucleotide promoter region surrounding the transcription start site of the human genome (hg38) of each gene was screened for transcription factor binding site (TFBS) regulatory motifs using the Bayesian regulatory site prediction algorithm MotEvo [55], which incorporates information from orthologous sequences in six other mammals and uses explicit models for the evolution of regulatory sites. The 190 regulatory motifs represent binding specificities of roughly 350 different human TFs. These were lifted to rn5 using liftOver [56]. The predicted number of functional TFBSs for each promoter was summed and used to perform Fisher Enrichment analysis between the genes up-regulated in NGF and NT3 conditions. The result of this analysis is presented in **Supplementary Table 3**.

### Analysis of Alternative cleavage and polyadenylation (APA)

In order to systematically investigate the changes in the poly(A) site usage between conditions we first scored the log2 proximal-to-distal poly(A) site ratio in each compartment or condition and each individual biological replicate by taking the log2 ratio of promoter-proximal and promoter-distal 3’ UTR isoform expression levels, hereafter called RUD. A score higher than 0 therefore indicates higher abundance of promoter-proximal 3’UTR isoform in the compartment or condition of interest. In the case of multiple promoter-distal 3’ UTR isoforms per transcript ID, a value was computed for each individual promoter-distal 3’ UTR isoform i.e. several values could be obtained per transcript ID family. We also computed the relative 3’ UTR usage between the promoter-proximal and the promoter-distal 3’ UTR isoform by calculating the ratios between the read count in the promoter-proximal and the sum of the read counts in the promoter-proximal and the promoter-distal 3’ UTR isoforms, hereafter called PUD.

In order to identify tandem 3’ UTR that show a marked change in the use of promoter proximal and distal poly(A) sites either between NGF and NT3 in the cell body or between cell body and axons in either NGF or NT3, we restricted our analysis on transcript ID families containing at least two 3’ UTR isoforms, and for which at least one promoter-distal 3’ UTR isoform was expressed in at least one of the compared conditions. In order to identify transcripts that show a marked change in the pA site usage between conditions, we scored the differences in proximal-to-distal poly(A) site usage using the following two scores:

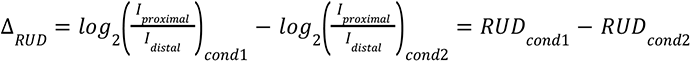

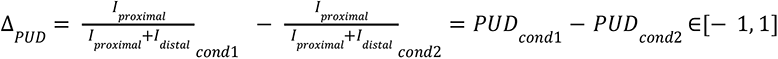

The statistical significance of the changes in proximal-to-distal poly(A) site ratio between conditions was assessed by Fisher’s exact count test using summed-up raw read counts of promoter-proximal versus promoter-distal 3’ UTR isoforms originating from either conditions (cell body versus axonal compartment or cell bodies of NGF versus NT3 conditions). We adjusted the P-Value controlling for False Discovery Rate (FDR) of 0.01. We restricted our analysis on Ensembl transcripts containing at least two 3’ UTRs generated by tandem polyadenylation expressed in either condition. Proximal shifts were then selected when Δ*_RUD_* ≤ – 1, Δ*_PUD_* ≤ – 15% and FDR<0.01; distal shifts were selected when selected when Δ*_RUD_* ≥ 1, Δ*_PUD_* ≥ 15% and FDR<0.01.

### Axonal localisation analysis

The ratio between gene’s abundance in the axons and in the cell body is often used to quantify mRNA axonal localisation [37, 57], however this metric correlates with the mRNA abundance in the cell body and the transcript length (**Supplementary Figs. 3C,D**). Consequently, such metrics fail to identify highly transported but lowly expressed transcripts, and similarly they are more likely to associate highly expressed transcript with high mRNA axonal localisation scores albeit low transport efficiency as compared to transcripts of similar expression level. Here we aimed to develop a novel metric that accurately infers the axonal transport efficiency and stability, irrespective of the transcript length and the expression in the cell body using a hierarchical Bayesian model procedure.

First, we looked at the global relationship between the normalised read count per 3’ UTR isoforms in the axonal compartment and their corresponding normalised read counts in the cell body compartments. We found that the average 3’ UTR isoform abundance level -average log2 expression-in the axonal compartments of either NGF or NT3 condition is best approximated by a combination of polynomial regression model of degree four of the abundance in the cell body completed and a linear model of the transcript length as revealed by the Akaike’s An Information Criterion from ANOVA analysis (**Supplementary Table 22** and **Supplementary Fig. 3E**). Given our goal to set out a metric of axonal localisation independent of the cell body read counts and the transcript length, we created 102 groups of 3’ UTR isoforms of fixed expression ranges in the cell body (18 bins from 2^3^ to 2^20^ nucleotides) and fixed ranges of transcript lengths (10 bins from 10^2^ to 10^4.5^ nucleotides). For each of these 102 groups, we generated a matrix of 10^4^ simulated draws of axonal read count values predicted using the fitted polynomial regression model of degree four, obtained from the total pool of transcripts, over a regularly interspaced grid of 100 possible transcript lengths -ranging from the minimal to the maximal transcript length of this specific group, and 100 possible cell body read count values -again ranging from the minimal to the maximal cell body read count of this specific group. We then extracted the average and variance over the 10^4^ axonal read count values for each of these 102 groups. Using these 102 pairs of averages and variances, we next fitted 102 negative binomial distributions to the 102 groups of axonal read counts by maximum likelihood (mle) using the fitdist function from the fitdistrplus R package [58]. These 102 posterior predictive distributions of axonal read counts could then serve to assess whether the observed axonal abundance of each individual 3’ UTR isoform 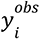 are consistent with the fitted models given their associated transcript length and cell body abundance. Here we propose that the more extreme the 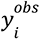, for a given 3’ UTR isoform of specific transcript length and cell body abundance, is on the histogram of simulated values 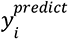 from the corresponding predictive distribution, the more likely the axonal localization of its corresponding 3’ UTR isoform has been actively regulated as opposed to the unspecific active transport which is expected to affect most transcripts detected in the axons. If 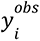 is in the lower tail, we expect this 3’ UTR isoform to be either restricted to the cell body or actively degraded in the axonal compartment; conversely if 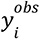 is in the higher tail of the histogram, we expect the 3’ UTR isoform to be either actively transported or stabilised in the axonal compartment. Thus for each 3’ UTR isoform *i*, we next computed the proportion of 10^4^ values, randomly generated from the posterior negative distribution associated with its corresponding transcript length and abundance in the cell body, that were smaller or larger than the observed axonal read count. In order to get a single value per transcript, hereafter called *Localisation Score*, we selected the smallest of these two values (probabilities to observe smaller or larger axonal read count for a given isoform), transformed it using the log10, and multiplied it by -1 when the latter was selected. Hence positive values are associated with active transport or stabilisation, while negative values are associated with cell body restriction or degradation. Notably while the ratios between gene’s abundance in the axons and in the cell body depend on cell body abundance (**Supplementary Fig. 3F**, *upper*) and transcript length (**Supplementary Fig. 3G**, *upper*), this is not the case anymore with the novel localisation score metric (**Supplementary Figs. 3F,G**, *lower*). This analysis has been restricted on the 30,450 3’ UTR isoforms detected in the axonal compartments of neurons either exposed to NGF or NT3. The axonal localisation scores for these 3’ UTR isoforms have been computed for NGF and NT3 conditions independently and are reported in **Supplementary Table 9**.

### Mapping and analysis of CLIP data

To identify RBPs that bind to 3’ UTR regions, we examined iCLIP data for 18 RBPs [59], and eCLIP data from K562 and HepG2 cells for 89 and 70 RBPs, respectively, available from ENCODE [13, 60]. In total we analysed Clip-seq data for 126 RBPs. Before mapping the reads, adapter sequences were removed using Cutadapt v1.9.dev1 and reads shorter than 18 nucleotides were dropped from the analysis. Reads were mapped with STAR v2.4.0i [61] to UCSC hg19/GRCh37 genome assembly. To quantify binding to individual loci, only uniquely mapping reads were used. The results were lifted to rn5 using liftOver [56].

#### RBP regulome underlying alternative polyadenylation

In order to identify positive and negative RBPs regulators of alternative polyadenylation in developing rat sympathetic neurons, we tested the association between RBP binding in defined regions along the 3’ UTR and the relative usage of the promoter-proximal 3’ UTR isoforms (PUD) in the cell body compartments of either NGF or NT3 conditions. In particular we used the Welch’s t-test to compare the distributions of the PUD between groups of isoforms exhibiting or not cross-link events in the following 30 defined regions surrounding the 3’ ends of the *promoter-proximal* 3’ UTR isoforms :{ [-3000:-2950],[-2750:-2700],[-2500:-2450],[-2250:-2200],[-2000:-1950],[-1750:-1700],[-1500:-1450],[-1400:-1350],[-1300:-1250],[-1200:-1150],[-1100:-1050],[-1000:950],[-900:-850],[-800:-750],[-700:-650],[-600:-550],[-500:-450],[-450:-400],[-400:-350],[-350:-300],[-300:-250],[-250:-200],[-200:-150],[-150:-100],[-100:-50],[-50,0],[0,+50],[+50:+100],[+100:+150]} for 126 RBPs with available CLIP-seq data in human cell clines, thereby obtaining 30 *P*-values per RBPs. These were then minus log10-transformed and multiplied by the sign of the difference in PUD between the group of isoforms exhibiting or not a cross-link event in the defined region along the 3’ UTR. Thus positive value indicates that RBP binding to regions surrounding the 3’ end of the *promoter-proximal* 3’ UTR promotes the usage of the promoter-proximal 3’ UTR isoforms, hence acting as positive regulators of the promoter-proximal and negative regulators of promoter-distal 3’ UTR isoform, while negative value indicates that RBP binding to regions surrounding the 3’ end of the promoter-proximal 3’ UTR promotes the usage of the promoter-distal, hence acting as negative regulators of the promoter-proximal and positive regulators of promoter-distal 3’ UTR isoforms. We repeated this analysis to compare the distributions of the PUD between groups of isoforms exhibiting or not cross-link events in the same 30 defined regions surrounding the 3’ ends of the *promoter-distal* 3’ UTR isoforms, therefore inspecting the regulatory potential of RBP through the binding of the distal 3’ UTR isoforms. For each of the 126 RBPs we therefore obtained a map of regulatory potential of the relative 3’ UTR usage along the short and long 3’ UTR.

#### RBP regulome underlying APA between NGF and NT3 and axonal remodelling

To identify RBPs regulators of APA between NGF and NT3, we performed Fisher count enrichment analysis to test for significant enrichment in RBP cross-link events in defined regions (see previous paragraph) around the 3’ end of either the promoter-proximal or the promoter-distal 3’ UTR isoform between the groups of isoforms exhibiting significant shifts in either groups and the total pool of 3’ UTR isoforms. This analysis is indeed more efficient in recovering candidate RBP regulators as compared to the Welch’s t-test comparing the distributions in Δ*_PUD_* between groups of isoforms exhibiting or not a cross-link event given the very low number of pairs of isoforms exhibit promoter-proximal or distal shifts in the cell bodies of NGF and NT3 treated neurons. We used a similar approach to identify candidate regulators of axonal remodelling, testing for significant enrichment in RBP cross-link events in defined regions around the 3’ end of the promoter-proximal 3’ UTR isoforms between the groups of promoter-proximal 3’ UTRs exhibiting axonal remodelling in either NGF or NT3 conditions as compared to the total pool of promoter-proximal 3’ UTR.

#### RBP regulome underlying axonal transport

To identify positive and negative RBPs regulators of axonal 3’ UTR isoform localisation and stability in developing rat sympathetic neurons, we tested the association between RBP binding in defined regions along the 3’ UTR and the localisation score in NGF or NT3 conditions. In particular we used the Welch’s t-test to compare the distributions of the localisation scores between groups of isoforms exhibiting or not cross-link events in 30 defined regions surrounding their 3’ ends (see above for detailed regions), for 126 RBPs with available CLIP-seq data in human cell lines, thereby obtaining 30 *P*-values per RBPs. These were then minus log10-transformed and multiplied by the sign of the difference in localisation scores between the group of isoforms exhibiting or not a cross-link event in the defined region along the 3’ UTR. Thus positive values indicates that RBP binding to regions surrounding the 3’ end of 3’ UTR isoform promotes axonal transport and/or stability, while negative values indicates that RBP binding to regions surrounding the 3’ end of 3’ UTR isoforms prevents axonal transport and/or promotes mRNA degradation.

### Gene Ontology (GO) enrichment analysis

GO analysis was performed by comparing pairs of gene lists using the Fisher test with the topGO Bioconductor package [62]. Only GO terms containing at least 10 annotated genes were considered. We applied a P-value threshold of 0.05. We manually filtered biologically relevant and statistically enriched GO by removing redundant GO terms and those applying to fewer than 5 genes in the gene lists.

### Statistical analyses

Data are expressed as averages ± SEM. t-test was used as indicated to test for statistical significance, which was placed at least P < 0.05 unless otherwise noted.

## Availability of data and materials

The accession number for the sequencing data used in this study is GSE160025. All the custom code required to infer 3’ end from sequencing data together with the GTF annotation file can be freely accessed on GitHub: https://github.com/RLuisier/my3UTRs.

## Ethics approval and consent to participate

Not applicable.

## Consent for publication

Not applicable.

## Competing interests

The authors declare that they have no competing interests.

## Funding

The work was supported by the Wellcome Trust Investigator Awards 103717/Z/14/Z and 217213/Z/19/Z (to A.R.), the MRC LMCB Core Grant MC/U12266B (to A.R), a Wellcome Trust Institutional Strategic Support Fund (to C.A.), and Idiap Research Institute (to R.L.)

## Acknowledgements

We thank all members of the Riccio and Luisier laboratories, as well as Pierre Klein for stimulating discussions and critical review of the data.

## Authors Contribution

R.L., C.A. and A.R. conceived and designed the study. C.A. performed the screen and all experiments presented in the study. R.L. designed the computational framework, performed data analyses and interpretation of results, designed the figures and derived the model. R. L. and A.R. wrote the manuscript with the critical input from C.A. All authors discussed the results and contributed to the final manuscript.

## Authors’ Twitter handles

Twitter handles: @RLuisier (Raphaëlle Luisier), @ar888 (Antonella Riccio).

## Tables

Supplementary tables S1-S23 can be accessed here.

Table S1 : Up-regulated genes in NGF treated cells compared to NT3

Table S2 : Up-regulated genes in NT3 treated cells compared to NGF

Table S3 : Transcription factor binding site enrichment analysis

Table S4 : Distal-to-proximal promoter 3’ UTR shifts in cell bodies of NGF treated cells compared to NT3

Table S5 : Distal-to-proximal promoter 3’ UTR shifts in cell bodies of NT3 treated cells compared to NGF

Table S6 : Description of the publicly available CLIP-sequencing data generated in human cell lines against 124 RBPs.

Table S7 : 17 positive regulators of polyadenylation

Table S8 : 27 negative regulators of the short 3’ UTR and positive regulators of the long 3’ UTR

Table S9 : Modelling of axonal localisation score

Table S10 : Over-transported transcripts in NGF-treated axons as compared to NT3-treated axons.

Table S11 : Over-transported transcripts in NT3-treated axons as compared to NGF-treated axons.

Table S12 : Positive RBP regulators of axonal localisation.

Table S13 : Negative RBP regulators of axonal localisation.

Table S14 : Promoter-proximal shifts in NGF-treated axons as compared to cell bodies

Table S15 : Promoter-proximal shifts in NT3-treated axons as compared to cell bodies

Table S16 : Promoter-distal shifts in NGF-treated axons as compared to cell bodies

Table S17 : Promoter-distal shifts in NT3-treated axons as compared to cell bodies

Table S18 : Candidate axonal remodeling in NGF

Table S19 : Candidate axonal remodeling in NT3

Table S20 : NGF RBP regulators of axonal remodeling

Table S21 : NT3 RBP regulators of axonal remodeling.

Table S22 : ANOVA linear model of axonal localisation.

Table S23: Primers and PCR conditions.

---

**Supplementary Figure 1.**
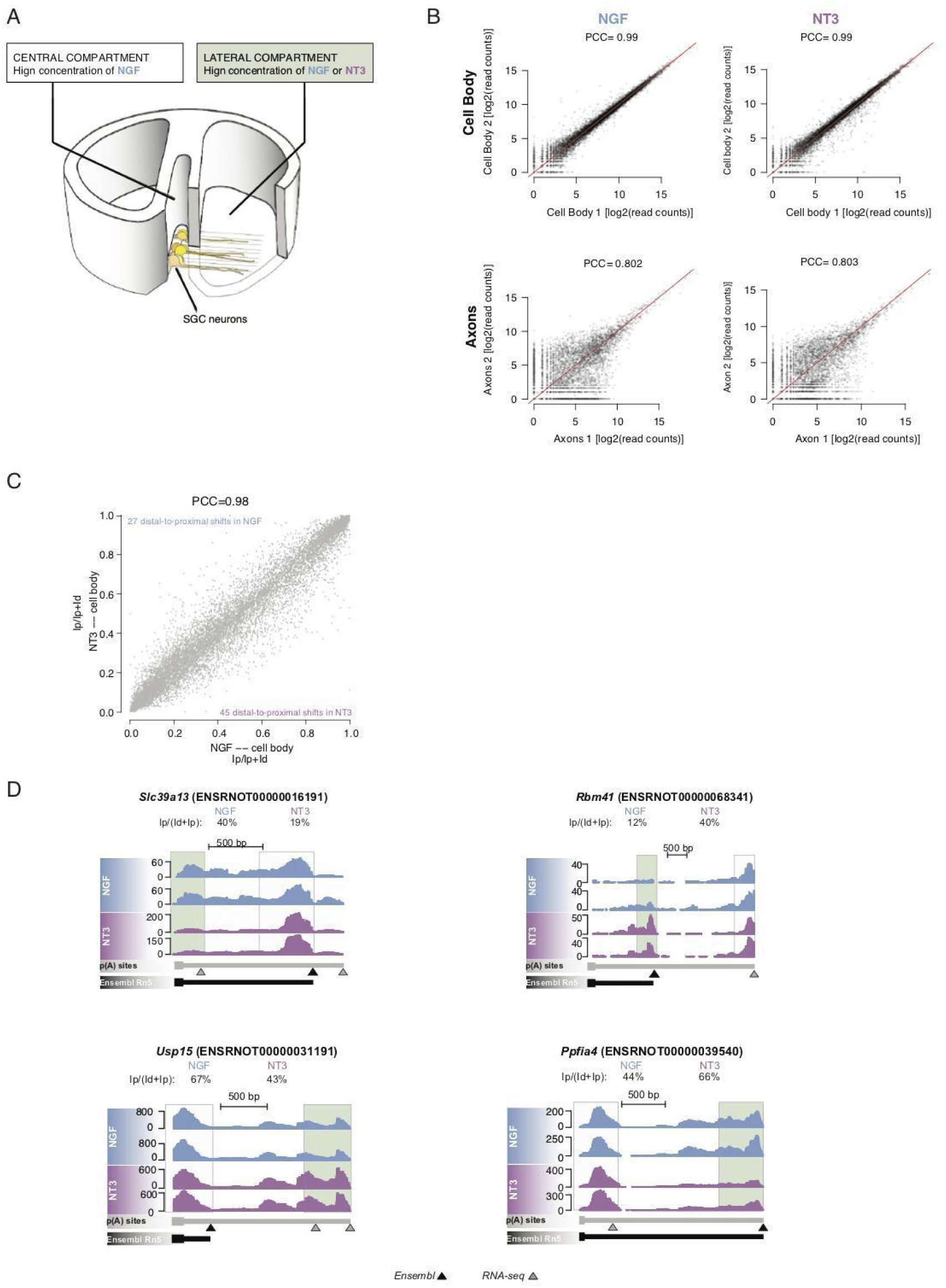
**(A)** Cell culture system used to isolate axonal or cell body mRNA from sympathetic neurons. **(B)** Scatterplot of the gene expression level between technical replicates. PCC=pearson correlation score. **(C)** Scatterplot of the relative usage of promoter-proximal and promoter-distal 3’ UTR isoform in cell bodies of neurons exposed to either NGF or NT3 (FDR < 0.01 between NGF and NT3; Fisher’s exact test). Proximal shifts in NGF compared to NT3 (blue); proximal shifts in NT3 compared to NGF (purple). **(D)** Genome browser view of representative transcripts showing a marked shift toward increase in promoter-proximal 3’ UTR usage (Slc39a13 and Rbm41), or in promoter-distal 3’ UTR usage (Ppfia4 and Usp15) in either condition.

**Supplementary Figure 2.**
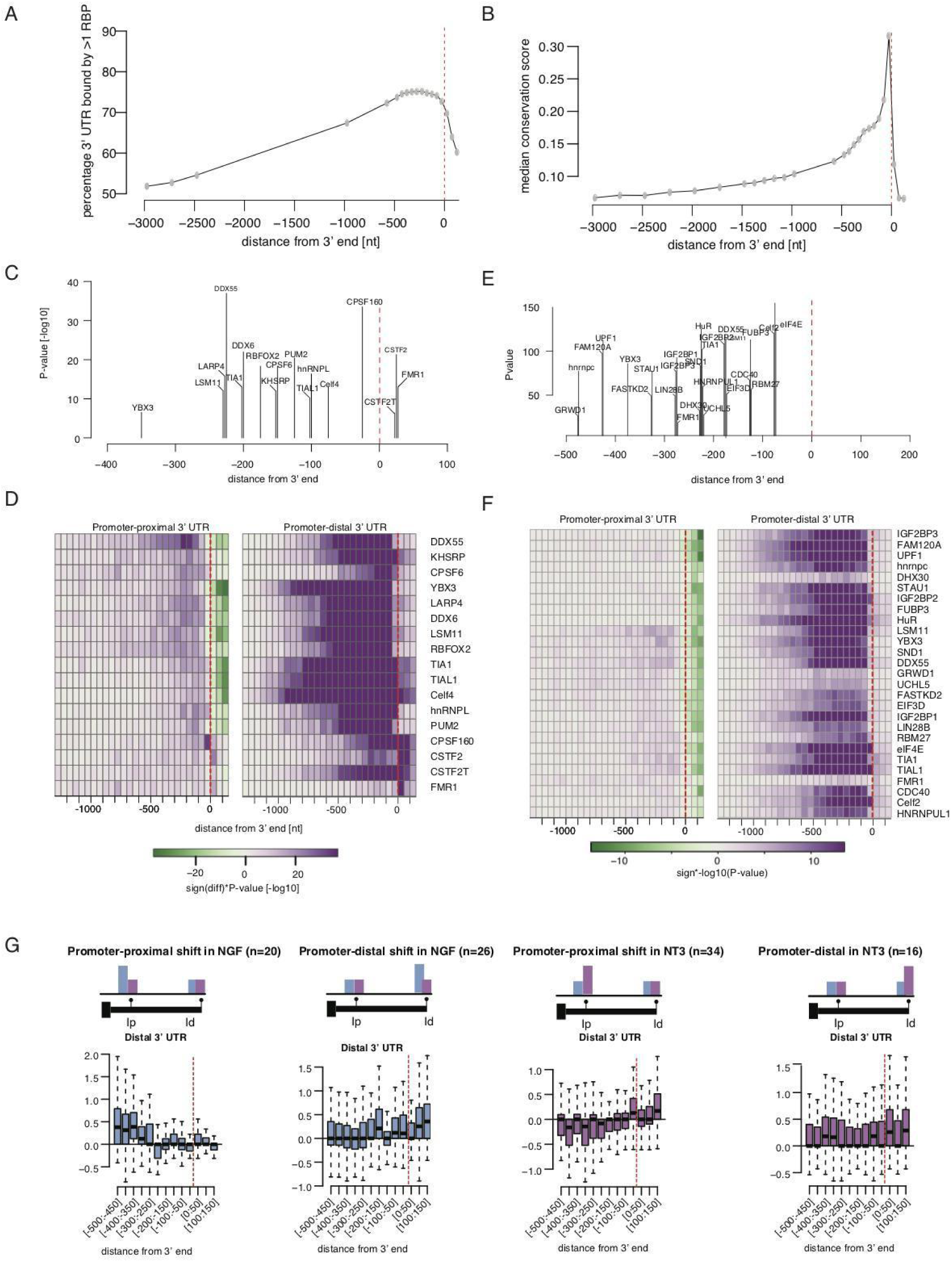
(A) Analysis of the percentage of 3’ UTR isoform exhibit cross-link event for at least one RBP as a function of the distance to the 3’ end. (B) Analysis of the nucleotide conservation score of 3’ UTR isoform in function of the distance to the 3’ end. **(C)** Position-dependent significance in positive association between cross-link events in defined regions along the short and long 3’ UTR for 17 identified positive regulators of polyadenylation. **(D)** Heatmap showing the extent of region-specific association between cross-link events for the individual 17 candidate positive regulators of polyadenylation and the relative usage of the proximal (*left*) or distal (*right*) 3’ UTR isoform. **(E)** Position-dependent significance in positive association between cross-link events in specific regions along the promoter-distal 3’ UTR for 27 identified positive regulators of long 3’ UTR and negative regulators of short 3’ UTR isoform. **(F)** Heatmap showing the extent of region-specific association between cross-link events for the individual 27 candidate negative regulators of promoter-proximal 3’ UTR isoforms and the relative usage of the proximal (*left*) or distal (*right*) 3’ UTR isoform. (**G**) Distribution of Fisher enrichment scores (-log10(P-value)) in RBP cross-link events along the long 3’ UTR isoforms of the pairs of isoforms exhibiting significant promoter-proximal shift or promoter-distal shift in NGF, and promoter-proximal shift or promoter-distal shift in NT3.

**Supplementary Figure 3.**
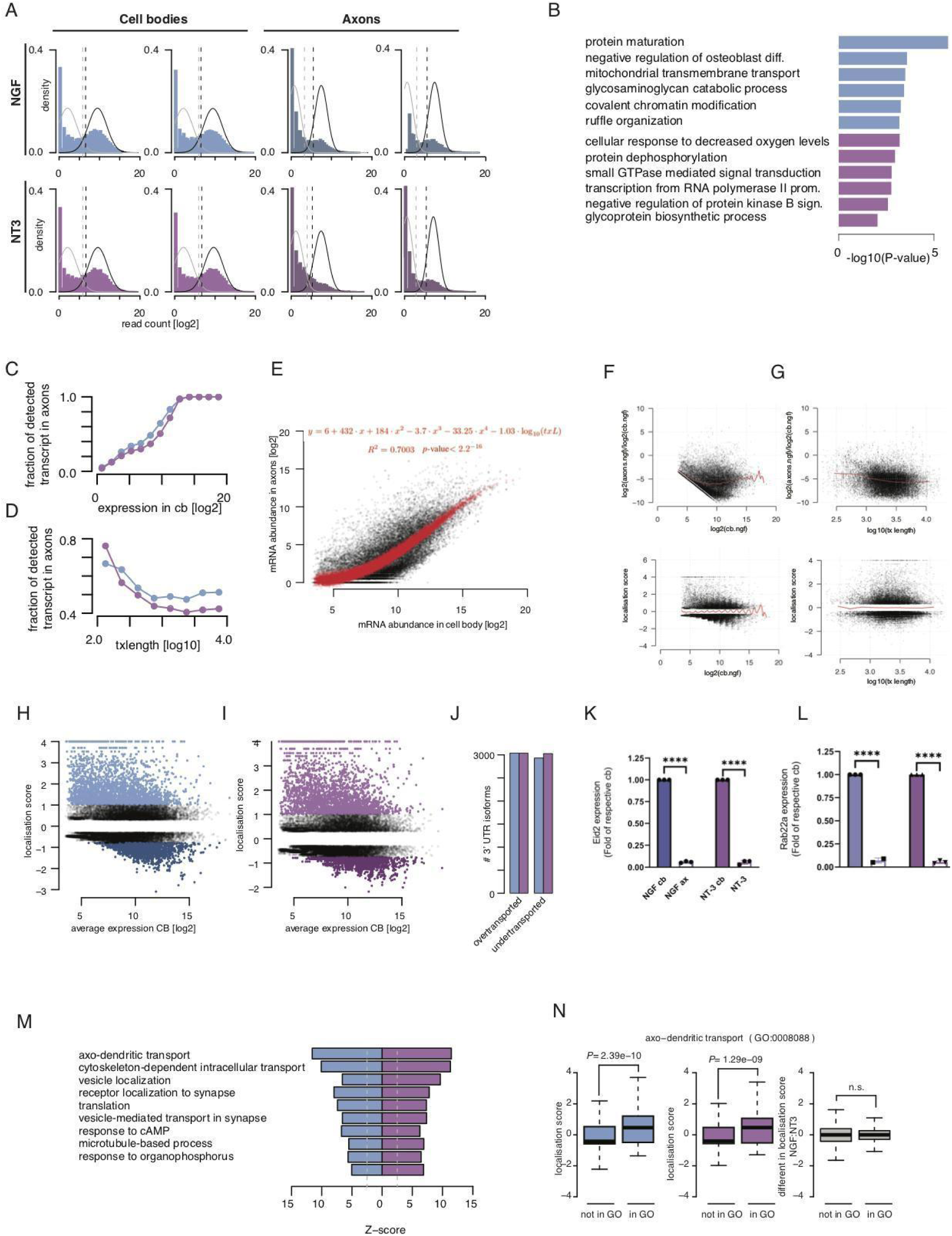
**(A)** Bimodal distribution of the log2-read count in cell bodies and distal axons samples. Dotted lines indicate the upper (stringent) and lower (loose) limits to discriminate the background gene expression level from the foreground. **(B)** GO biological pathway enrichment for genes detected in the distal axons of NGF (blue) or NT3 (purple) culture conditions only. **(C)** Fraction of detected transcripts in the axonal compartment and mRNA abundance in the cell body (log2). **(D)** Fraction of detected transcripts in the axonal compartment and transcript length (log10). **(E)** Relationship between the mRNA abundance in cell bodies (log2) and in distal axons (log2). A linear model is fitted that predicts the axonal mRNA abundance in function of the transcript length (log10) and of a polynomial of degree 4 of the abundance in the cell bodies (red equation). **(F)** Average mRNA abundance in the cell bodies (log2) and the ratios between the axonal and cell body mRNA abundance (log2; *upper*) or as compared with the localisation scores (*lower*). Red lines = average ratios (*upper*) or localisation scores (*lower*) per range of cell body abundance. **(G)** Transcript length (log10) and the ratios between the axonal and cell body mRNA abundance (log2; *upper*) or as compared with the localisation scores (*lower*). Red lines = average ratios (*upper*) or localisation scores (*lower*) per range of cell body abundance. **(H, I)** Localisation score and average mRNA abundance in the cell bodies of NGF- (J) or NT3-treated neurons (K). Dark dots=under-transported and light dots=over-transported. **(J)** Number of 3’ UTR isoforms exhibiting excessive axonal localisation (over-transported) as well as restricted to the cell bodies (under-transported) in either NGF or NT3 culture conditions. **(K)** Eid2 mRNA abundance in cell bodies and axons as measured by RT-qPCR (unpaired t-test, n=3, ****p<0.0001). **(L)** Rab22a mRNA abundance in cell bodies and axons as measured by RT-qPCR (unpaired t-test, n=2 or 3 as indicated, ****p<0.0001). **(M)** Standardized scores (Z-score) quantifying the excess in axonal localisation in NGF and NT3 for the top ten GO biological pathways. **(N)** Distributions for the background genes and the genes belonging to the axo-dendritic transport biological pathway of the localisation scores in NGF (*right*), the localisation scores NT3 (*center*), and the differences in localisation scores between the two conditions (*right*). (Welch’s t-test assessing the significant difference between the mean localisation scores. Boxplots display the five number summary of median, lower and upper quartiles, minimum and maximum values).

**Supplementary Figure 4.**
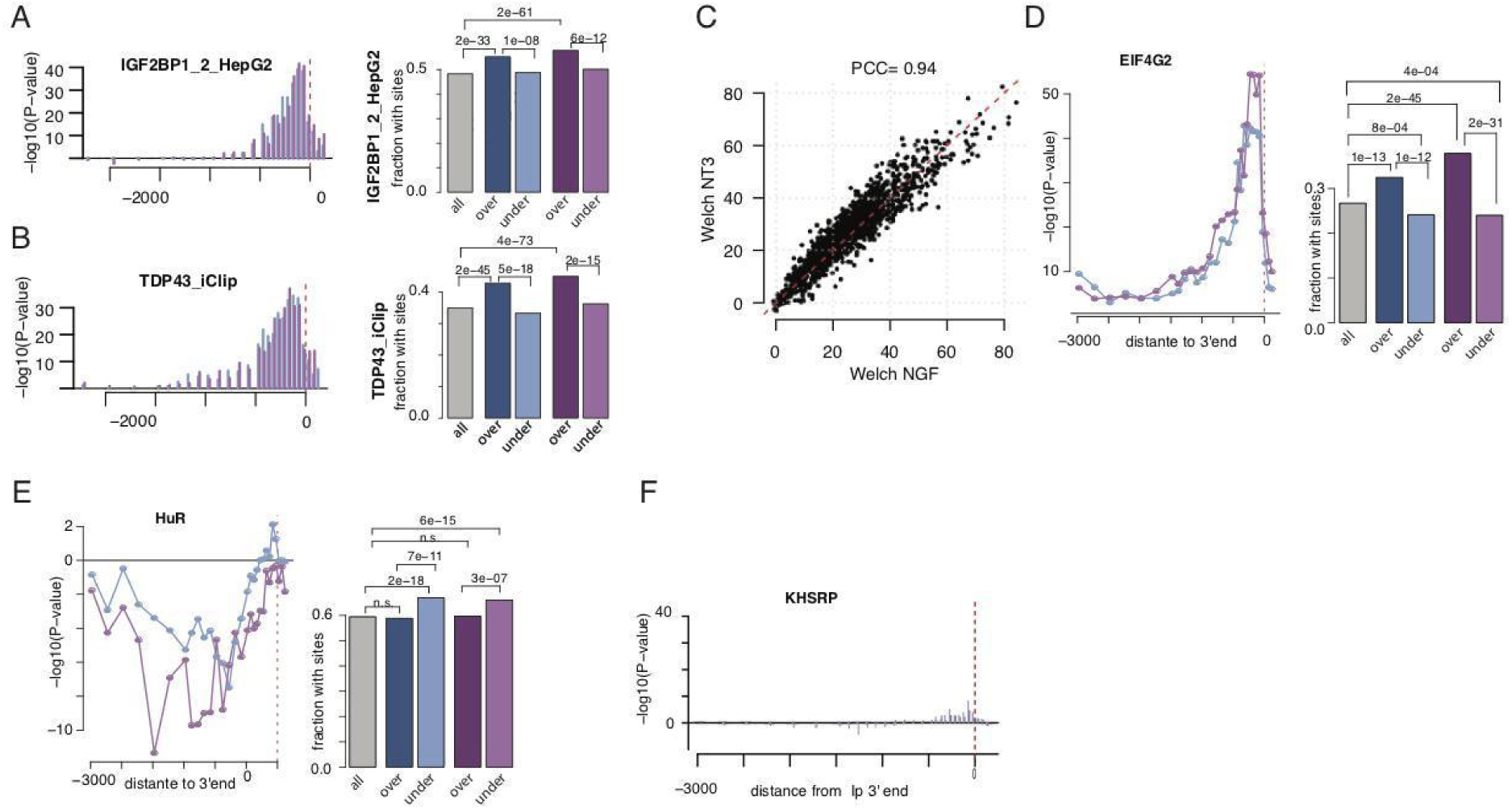
**(A)** Significant association between IGF2BP1 cross-link event and axonal localisation in defined regions along the 3’ UTR *(left*). Blue = NGF; purple = NT3. (*Right*) Barplots showing the fraction of [-250:-50] nucleotide regions upstream the 3’ ends exhibiting IGF2BP1 cross-link event in the full set of detected transcript (grey bar), and the pools of over- and under-transported transcripts in NGF (blue bars) and NT3 (purple bars), respectively. (Fisher enrichment test). **(B)** Same as (A) for TDP43. **(C)** Scatterplot of the extent of significance association between the 126 RBPs cross-link events in the [-250:-50] nt region upstream the 3’ end and the axonal localisation in NGF and NT3. **(D, E)** Same as (A) for HuR and KHSRP, respectively.

**Supplementary Figure 5.**
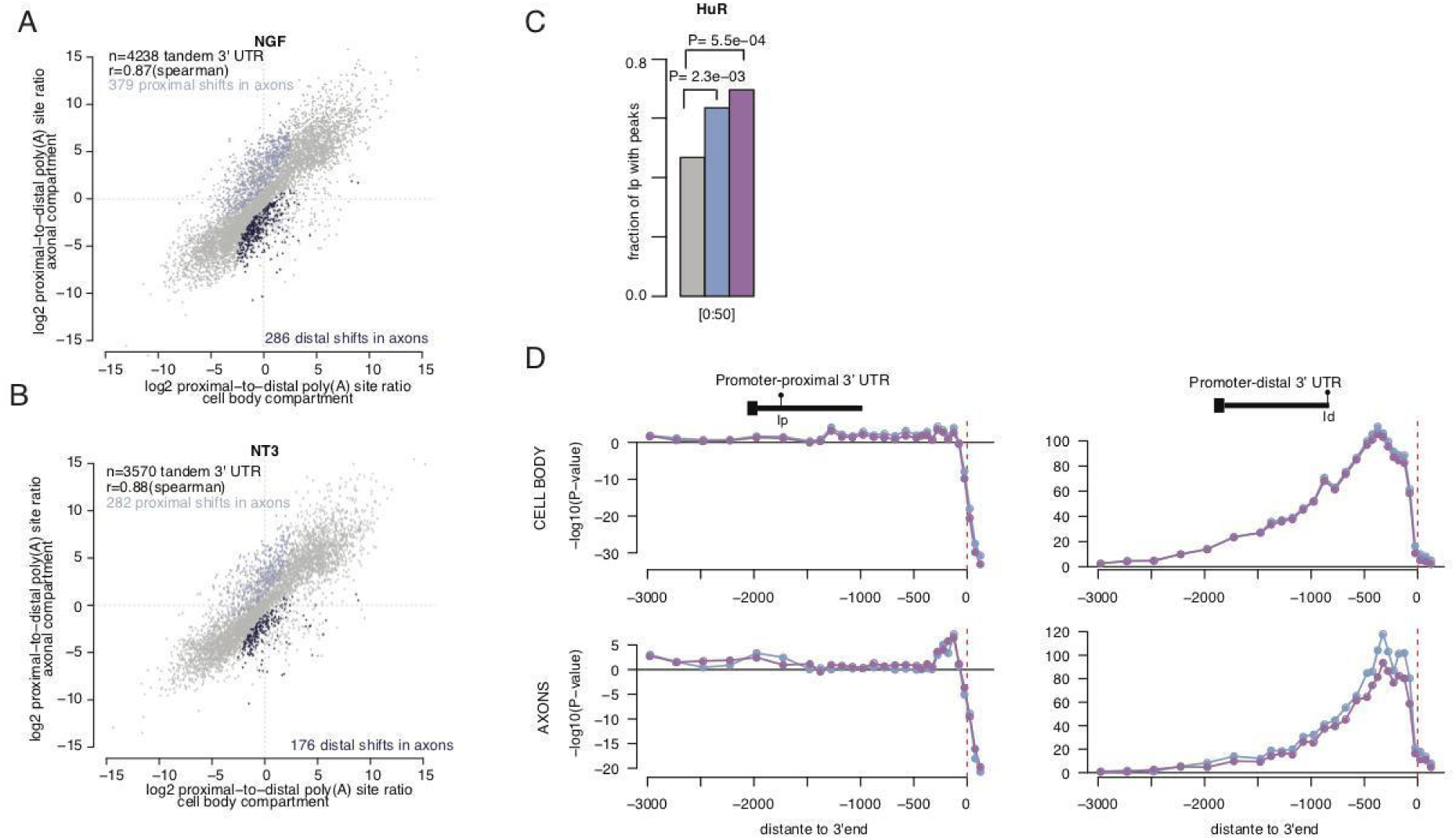
**(A)** Scatterplot of the relative usage of promoter-proximal and promoter-distal 3’ UTR isoforms sites in cell bodies and axons of NGF-treated condition (FDR < 0.01 between cell body and axonal compartment; Fisher’s exact test). Distal shifts in axons compared with cell body (dark blue); proximal shifts in axons compared with cell body (light blue). **(B)** Same as (F) for NT3-treated condition. **(C)** Fraction of promoter-proximal 3’ UTR isoforms exhibiting cross-link events for HuR in [0:50] nucleotide region down-stream of the cleavage site. Gray bar = all promoter-proximal 3’ UTR; blue bars = 80 candidates of axonal remodeling in NGF; purple bars = 60 candidates of axonal remodeling in NT3. (**D**) Scatterplot of the extent of significant association between UPF1 cross-link event in defined regions along the 3’ UTR and the relative usage of the promoter-proximal 3’ UTR (*left*) or the promoter-distal 3’ UTR (*right*) in the cell body (*upper*) or the axons (*lower*) . Blue = NGF; purple = NT3.

## Notes

https://docs.google.com/spreadsheets/d/1j41gYIQRUBWQgzSmyuemJLywWBChFeCsK3z9q6kopTI/edit?usp=sharing

### Competing Interest Statement

The authors have declared no competing interest.

